# Long-term vegetation changes in species-rich *Nardus* grasslands of Central Germany indicate eutrophication, recovery from acidification and management change as main drivers

**DOI:** 10.1101/543512

**Authors:** Cord Peppler-Lisbach, Nils Stanik, Natali Könitz, Gert Rosenthal

## Abstract

**Questions:** Which trends and patterns of community change occurred in *Nardus* grasslands over the past decades in parts of the Continental biogeographic region in Germany? Are patterns and trends similar at local and regional scale? Do impacts of environmental changes on *Nardus* grasslands in Central Europe correspond to those identified in the European Atlantic biogeographic region?

**Location:** East Hesse Highlands, Germany

**Methods:** In 2012-2015, we resurveyed vegetation relevés on quasi-permanent plots initially surveyed between 1971 and 1987 and re-measured soil parameters. We tested for differences in species frequency and cover, mean Ellenberg indicator values, species richness and soil variables. Nitrogen and sulphur deposition data were analysed to evaluate effects of atmospheric pollutants. We used regression analyses and redundancy analyses to identify environmental drivers responsible for changes in species composition.

**Results:** A regional scale, we found significant increases in soil pH, Ellenberg R and N values, plant nutrient indicators, forbs, species of agricultural and fallow grasslands. C:N ratio, *Nardus* grassland specialists, low-nutrient indicators and graminoids declined. Changes in species composition are related to changes in pH and management. There was a strong decrease in sulphur and a moderate increase in nitrogen deposition, whose local scale pattern did not correlate with changes in soil parameters. However, there was a statistical effect of local NHy changes on species composition.

**Conclusion:** The findings indicate significant overall eutrophication, a trend towards less acidic conditions, insufficient management and abandonment, which are widely consistent across study areas and correspond to recent reports of vegetation changes and recovery from acidification in the Atlantic biogeographic region. We assume reduction in sulphur deposition during recent decades to be a major driver of these changes combined with increased nitrogen deposition and reduced management intensity. This suggests a large-scale validity of processes triggering changes in *Nardus* grasslands of Western and Central Europe.

**Nomenclature:** The nomenclature follows the German taxonomic reference list (GermanSL version 1.3) of Jansen and Dengler (2008).

## Introduction

Semi-natural grasslands are of high importance for human well-being by providing important ecosystem services and high biodiversity (Dengler, Janisova, Toeroek, & Wellstein, 2014; Hejcman, Hejcmanova, Pavlu, & Benes, 2013). They are, however, under threat by effects of global change, e.g. land-use change, nitrogen deposition and climate change (Sala et al., 2000). Recent decades brought evidence for the important role of atmospheric depositions on grassland biodiversity, mainly of nitrogen (N) and sulphur (S) (Morecroft et al., 2009). Consequences for European semi-natural grasslands are the loss of species diversity, a change in species composition and a decline of ecosystem functions (Bobbink et al., 2010; Phoenix et al., 2012; Stevens, Dise, Mountford, & Gowing, 2004).

*Nardus* grasslands are typical species-rich semi-natural grasslands on strong to moderate acid soils in large parts of temperate Europe. They are often also referred to as or included in the type ‘acid grasslands’ (e.g. Damgaard et al., 2011; Stevens, Duprè, Gaudnik et al., 2011). In the European context, they are classified as the priority natural habitat (“Species-rich *Nardus* grasslands, on siliceous substrates in mountain areas”, H6230*) of the EU Habitats Directive (Directive 92/43/EEC, European Council, 1992). *Nardus* grasslands harbour a considerable number of endangered species. For Germany, 19 species from the Red List are characteristic for *Nardus* grasslands and the habitat hosts a high number of threatened animal taxa such as meadow-breeding birds, butterflies and wild bees (Peppler-Lisbach & Petersen, 20010; Schwabe et al., 2019). Historically connected to low-input land-use practices, they are important constituents of pre-industrial land-use systems and thus part of the European cultural heritage (Galvánek & Janák, 2008). Therefore, conservation efforts of the European community aim to protect these grasslands in a favourable conservation status.

Indicated by their legal protection status, *Nardus* grasslands are highly endangered due to global change drivers. Among these drivers, mainly abandonment and land-use intensification have triggered the decline of *Nardus* grasslands in Central Europe since the late 19^th^ century (Leuschner & Ellenberg, 2017). Moreover, their preference for poorly buffered, nutrient-poor soils makes them particularly vulnerable to processes of eutrophication and acidification (Dupré et al., 2010; Helsen, Ceulemans, Stevens, & Honnay, 2014). This applies also to Atlantic heathlands (Bobbink, Hornung, & Roelofs, 1998; Southon, Field, Caporn, Britton, & Power, 2013), to which *Nardus* grasslands are floristically related. The goal of this study is to investigate long-term changes in *Nardus* grassland in the Continental biogeographic region and to show the extent to which these drivers hinder the conservation of this habitat type.

Eutrophication summarises effects of nutrient enrichment, mainly of nitrogen or phosphorous, and can be attributed to both agricultural fertilisation and atmospheric deposition (Bobbink et al., 2010; Ceulemans, Merckx, Hens, & Honnay, 2013). It results in species losses due to competitive exclusion of light-demanding, low-competitive species and other changes in species and functional composition (Bobbink & Hicks, 2014; Helsen et al., 2014). Acidification is mainly driven by atmospheric deposition of N (NHy, NOx) and S (SOx) but is also modified by local factors like soil characteristics (Roy et al., 2014). It results in decreasing pH values, leaching of base cations, Al^3+^ mobilisation, and higher ratios of Al:Ca and NH_4_:NO_3_ (Kleijn, Bekker, Bobbink, de Graaf, & Roelofs, 2008; Ross et al., 2012; Stevens, Dise, & Gowing, 2009). These effects are considered to be responsible for the decline and regional extinction of small-growing *Nardus* grassland and heathland species in the Netherlands (de Graaf, Bobbink, Roelofs, & Verbeek, 1998; Fennema, 1991). Additionally, soil acidification may lead to reduced nutrient availability preventing the effects of N deposition from being effective (Stevens, Thompson, Grime, Long, & Gowing, 2010).

Studies about effects of N deposition on acid grasslands are predominantly from the Atlantic biogeographic region of Europe. Commonly reported effects are a decrease in total species richness, a decline in typical acid grassland or heathland species adapted to low nutrient availability, an increase in graminoid cover but a decrease in graminoid richness (Damgaard et al., 2011; Field et al., 2014; Payne et al., 2017; Stevens, Duprè et al., 2010). Moreover, decreases in forb and bryophyte cover and richness were found (Stevens, Dise, Gowing, & Mountford, 2006). Ellenberg N indicator values increased with increasing N deposition (Henrys et al., 2011; Pakeman et al., 2016), whereas R indicator values decreased due to N deposition-driven acidification (Maskell, Smart, Bullock, Thompson, & Stevens, 2010). Regarding environmental factors, soil pH was negatively correlated to N deposition rates, whereas C:N ratio showed a significant positive relationship with N deposition (Stevens et al., 2006; 2011).

While S deposition, after peaking in the 1980/90s, decreased considerably over the last two decades in many European regions (Kirk, Bellamy, & Murray Lark, 2010; Morecroft et al., 2009; Teufel, Gauger, & Braun, 1994), N deposition rates are still on a high level (Dentener et al., 2006), especially for NHy (Gauger, Nagel, Schlutow, & Scheuchner, 2013). Morecroft et al. (2009) and McGovern, Evans, Dennis, Walmsley, and McDonald (2011) associated decreasing SOx deposition with increasing soil pH values but did not observe any vegetation recovery from acidification. Hence, Stevens, Payne, Kimberley, and Smart (2016) assumed longer time scales for recovery signals of the vegetation. Predictions of Stevens, Duprè, Dorland et al. (2011) about impacts of N deposition on soil (pH, NHy:NOx ratio) and species shifts (from stress-tolerant species to more competitive species) were recently shown in acid grasslands (Mitchell et al., 2018; Rose et al., 2016).

Effects of land-use changes contribute additionally to long-term N and S deposition-driven vegetation changes in semi-natural grasslands (Humbert, Dwyer, Andrey, & Arlettaz, 2016). For example, infrequent mowing and low or reduced grazing intensity promotes dwarf shrubs and tall tussock grass species due to lower disturbance intensity. Simultaneously, characteristic small-growing *Nardus* grassland specialists are outcompeted and specific habitat structures, like increased litter or moss cover, are facilitated (Arens & Neff, 1997; Armstrong, Grant, Common, & Beattie, 1997). Contrastingly, an intensive management of *Nardus* grasslands by increased livestock grazing, early and more frequent mowing favours highly competitive common grassland species and the homogenisation of community structures (Mariotte, Buttler, Kohler, Gilgen, & Spiegelberger, 2013).

This study is based on local scale investigations of Peppler-Lisbach and Könitz (2017), who detected long-term changes in *Nardus* grasslands in a small study area in Central Germany (Fulda-Werra-Bergland). They revealed eutrophication effects, but no acidification during the time span of the resurvey and rather found indications of recovery from acidification. For the present study, we aim to upscale findings from local scale to regional scale by considering and adding data from a second resurvey study area from the Continental biogeographic region. Moreover, we target to detect common trends and spatial patterns between changes in deposition, soil parameters and species composition to gain insights into causal relationships between spatially explicit changes in N and S deposition regimes and vegetation change via environmental factors. Accordingly, we hypothesise that there have been considerable changes in N- and S-deposition rates between initial survey and resurvey in both study areas. As a result, we expect that changes in ecosystem-relevant depositions triggered changes in soil conditions, which can be identified exemplarily by a decrease in C:N ratio and an increase in soil pH as indications for eutrophication and recovery from acidification, respectively. Hence, we assume that changes in soil chemical drivers and possible changes in management induced significant changes in vegetation composition of *Nardus* grasslands not only at local but also at regional scale. This would confirm recently observed long-term vegetation changes to have a consistent pattern at regional scale and would support a cross-regional consistent recovery trend from acidification as reported by Rose et al. (2016) and Mitchell et al. (2018) from the Atlantic biogeographic region.

## Material and methods

### Study areas

The study areas (local scale: Fulda-Werra-Bergland, FWB, and Rhön Mountains, RHN) are parts of the East Hesse Highlands (regional scale) in the Central German low mountain range and about 100 km apart from each other. Both areas have a geological base predominantly built of Triassic sandstone, which is locally (FWB) or dominantly (RHN) covered by tertiary basalt. The altitudes of the study plots vary from 230 to 720 m a.s.l. (FWB) and 570 to 940 m a.s.l. (RHN). The climate is of a sub-oceanic character with a mean annual precipitation of 650-1000 mm and mean annual air temperature of 5-9°C (Bohn, Korneck, & Meisel, 1996; Klink, 1969).

### Data collection

#### Vegetation surveys

The study is based on a resurvey of vegetation relevés of *Nardus* grasslands (*Nardetalia strictae, Violion caninae*) (Peppler-Lisbach & Petersen, 2001). Plots (97 in total) were initially surveyed in 1971 by Borstel (1974) (*n* = 10, RHN) and in 1986/87 (*n* = 87, RHN: 27, FWB: 60) by Peppler (1992), respectively. The plots in each study area were resurveyed in 2012 (FWB) and 2014/15 (RHN). Plot sizes were taken from the initial surveys and varied between 6 and 50 m² (mean 23.6 m², sd 8.2). To eliminate influences of direct fertilisation and succession, we excluded resurvey plots in RHN, which experienced agricultural intensification, and plots that experienced advanced undisturbed secondary succession since the initial survey (i.e. shrub/tree layer > 60% cover). Hence, all remaining plots either are within protected areas or managed according to agri-environment schemes without fertilisers. Species cover/abundance values were harmonised on the standard Braun-Blanquet scale (r, +, 1-5). The plots can be classified as “quasi-permanent plots” (Kapfer et al., 2017), as they were not permanently marked but could be relocated using precise hand-drawn maps and geographical coordinates; following the recommendations of Kapfer et al. (2017) to reduce the inherent error in this source of resurveyed data.

#### Soil sampling and analysis

Mixed soil samples of the upper 0-10 cm (auger diameter 5 cm) were collected during the resurvey. Samples were thoroughly mixed to ensure homogeneity and were sieved to < 2mm for further processing. Soil pH was measured electrometrically in deionised water (relevés of Peppler 1992) and 1 N KCl solution (relevés of Borstel 1974). Total C and N content of all soil samples to receive the C:N ratio (CN) was analysed using a CN element analyser (vario MAX CHN – Elementar Analysensysteme GmbH and Flash EA 2000 – Thermo Fischer Scientific; Germany).

#### Data on sulphur and nitrogen deposition

Deposition rates for both study areas were extracted for each plot from national modelling studies of airborne N (NHy, NOx, Ntotal) and S (SOx) pollution between 1987 and 2007 with a spatial resolution of 1 x 1 km (Gauger et al., 2002; Gauger et al., 2008; Gauger, 2010; Gauger, Kölbe, & Anshelm, 2000). Our plots were situated within 40 different grid cells (FWB: 21, RHN: 19). These modelled data are the only nationwide available data about N and S deposition in Germany covering both study areas in the period of comparison. Earlier modelled or measured data, to get cumulative deposition rates prior to the initial survey (e.g. Mitchell et al., 2018), were not available.

### Data analysis

Vegetation relevés were taxonomically harmonised. In doing so, some taxa had to be merged to aggregates (*Alchemilla vulgaris* agg., *Festuca ovina* agg. [including the character species *F. filiformis*] and *Ranunculus polyanthemos* agg.). Species were assigned to species groups according to three criteria (Supplementary information S1). Sociological groups based on their occurrence in certain syntaxa: character species (C, *Nardetalia* specialists in open habitats including heathland species according to Peppler-Lisbach & Petersen, 2001), other low-productive grassland species (D, species of anthropo-zoogenic heathlands/grassland according to Ellenberg, 1992 with an Ellenberg N indicator value of < 4), agricultural grassland species (G, species of anthropo-zoogenic heathlands/grasslands and an Ellenberg N indicator value of ≥ 4) and fallow species (F, species of forests, forest clearings and fringes according to Ellenberg, 1992, including trees and shrubs). Functional groups consist of graminoids (Gr, Poaceace, Cyperaceae and Juncaceae) and forbs (Fo, all other herbaceous, non-graminoid species). Ecological groups were defined according to species Ellenberg indicator values for soil reaction (R) and nitrogen (N): nutrient indicators (NI, N > 5), basiphytic low-nutrient indicators (bLI, N < 4 and R > 5) and acidophytic low-nutrient indicators (aLI, N < 4 and R < 4). For each sociological and ecological species groups, species numbers and cumulative cover were calculated for all of the 194 old and resurveyed relevés. For functional groups, we calculated the proportional richness and cover in relation to all species, trees and shrubs excluded (pGr and pFo, respectively) and the Gr:Fo ratio. For quantitative analyses of species cover values, original Braun-Blanquet cover codes were transformed to percentage values and subsequently square-root transformed. For all relevés, cover-weighted and unweighted (p/a) mean indicator values were calculated.

We derived the differences for all variables (v, i.e. species numbers, cumulative cover, environmental variables) between the initial survey (t1) and the resurvey (t2) as Δv = v(t2) – v(t1). To quantify changes in deposition rates, we averaged deposition rates for the modelling period 1987-89 (representing t1) and 2005-2007 (t2). ΔSOx, ΔNHy and ΔNOx were calculated accordingly. General trends in Δv were tested by one sample Wilcoxon tests (equivalent to paired two-sample Wilcoxon test, package “exactRankTest”, Hothorn & Hornik, 2017). To test for differences between study areas, we used two-sample exact Wilcoxon tests on Δv and study area as the grouping variable. Changes in soil variables, management, richness and species groups were tested for the plot-based dataset with 80 ≤ n ≤ 97, depending on data availability. To eliminate pseudo-replication in deposition data, we tested changes in deposition rates on a reduced dataset of the 40 different grid cells.

To test for effects of environmental drivers and study area, we calculated linear regression models for Δv as the respective dependent variable and differences in soil variables (ΔpH, ΔCN), changes in management (ΔM, three categories: xF [continuously or recently fallow, reference category], FM [t1: fallow, t2: managed], MM [t1: managed, t2: managed]), altitude (a.s.l.) and study area as predictor variables. Dependent variables were Box-Cox transformed prior to the analyses to improve normality. Variable selection was done by stepwise selection based on Bayes Information Criterion.

To analyse effects of changes in N and S deposition on species composition, we pooled plot values (vegetation data, soil variables, management) per grid cell to account for pseudo-replication caused by plots situated within the same grid cell of the deposition dataset (*n* = 40). As pooled values per grid cell, we used means (Δcover, ΔpH, ΔCN), medians (Δp/a) or the mode category (ΔM). Median Δp/a-values of 0.5/-0.5 were assigned to 1 and −1, respectively, to re-transform data to integer values.

As complemental approach to test for effects of drivers (ΔpH, ΔCN, ΔM, ΔSOx, ΔNHy, ΔNOx) and study area on the entire species composition, we employed redundancy analyses (RDA) of species differences (Δsqrt(cover) or Δp/a) and tested with permutation tests (R package “vegan”, Oksanen et al., 2015). These analyses were performed at plot level (*n* = 80) and when including deposition changes with the pooled dataset (*n* = 40). For deposition variables, we conducted single variable RDAs with study area as first constraining variable, the relating deposition change variable as second constraining variable and additionally their interaction with study area. Study area was taken as the first constraining variable to partial out local differences in species composition that could have been caused by other study area-specific factors.

For all statistical analyses including soil and structural variables, the respective number of observations varied with the availability of reference data and can be found in the result tables. All statistical analyses were conducted with the statistical software R 3.6.1 (R Core Team, 2017).

### Results

#### Changes in management

Contrary to the general high number of abandoned plots at the initial survey with 58 (i.e. 60 %) fallow plots, most plots (80, i.e. 82 %) were managed at the time of the resurvey (Table 1). Only three plots in the RHN and none in the FWB had been abandoned after the first survey. Concerning management type, mowing was realised in 77 % and grazing in 33% of all managed plots at the initial survey. This proportion changed due to an increase of grazed plots to 63 % and 37 % in 2012-2015. At that time, plots were managed either by late mowing (August or September) or by low-intensity grazing. Unfortunately, there were no consistent information available about the management of the plots between the two surveys and, in particular, about exact dates of re-introduced management on initially fallow plots.

**Table 1.** Management changes between time periods and study areas (FWB: Fulda-Werra-Bergland, RHN: Rhön Mountains).

| Management |  | Management change category | Study area |  |  |
| --- | --- | --- | --- | --- | --- |
| 1971-1987 | 2012-2015 |  | FWB | RHN | *n* |
| fallow | fallow | xF | 13 | 1 | 17 |
| mown | fallow | xF | 0 | 2 |  |
| grazed | fallow | xF | 0 | 1 |  |
| fallow | mulched | FM | 1 | 0 | 44 |
| fallow | mown | FM | 15 | 4 |  |
| fallow | grazed | FM | 21 | 3 |  |
| mown | mown | MM | 6 | 19 | 36 |
| mown | grazed | MM | 1 | 2 |  |
| grazed | mown | MM | 2 | 3 |  |
| grazed | grazed | MM | 1 | 2 |  |

Management change (ΔM) was translated into three categories: 17 plots were categorised as xF (i.e. fallow in 2012-2015), 36 as MM (managed in both periods) and 44 as FM (fallow-managed). The ΔM categories had an uneven altitudinal distribution (ANOVA: F_(2,94)_ = 13.18, P < 0.001), with MM concentrated at high altitudes (mean 732 m, sd 172 m), FM at intermediate altitudes (mean 558 m, sd 143 m) and xF at low altitudes (mean 530 m, sd 223 m). However, there was only a significant difference between MM and the other two ΔM classes (Scheffé test: P < 0.001).

#### Changes in sulphur and nitrogen depositions

At regional scale, there was a drastic decline in SOx deposition between 1987 and 2007 in both study areas (Table 2, Supplementary information S2), while NHy and Ntotal increased at the same time, albeit less pronounced. Contrary to NHy, NOx decreased slightly over time with no significant change in FWB. Although decreases in SOx and increases in NHy were significant in both study areas, there were marked differences in quantity. Reduction in SOx was higher in RHN, whereas increase in NHy was higher in FWB. The NHy:NOx ratio increased considerably in both study areas with no difference across study areas. Hence, the net increase in Ntotal for both study areas can be attributed solely to the increase in NHy. Changes in deposition rates of all components showed highly significant negative correlation with altitude (ΔSOx: r_p_ = −0.92, ΔNHy: r_p_ = −0.67, ΔNOx: r_p_ = −0.89 and ΔNtotal: r_p_ = −0.82, all P < 0.001). Thus, within the resurvey period there were stronger declines of deposition rates in lower altitudes than in higher altitudes. ΔNHy:NOx was however not correlated with altitude (r_p_ = 0.10, P = 0.56).

**Table 2.** Deposition rates of SOx, NHy and NOx at regional (overall) and local scale (FWB: Fulda-Werra-Bergland, RHN: Rhön Mountains): A) Means for 1987 (t1) and 2007 (t2) and B) Results of exact Wilcox-Tests: One sample tests on ΔSOx, ΔNHy, ΔNOx and FWB:RHN: two sample tests on differences between study areas. ↑ or ↓ indicate trends of change, while — indicates no change. Significance levels are indicated by * P < 0.05, ** P < 0.01, *** P < 0.001, ns: not significant. *n* = number of plots (overall/FWB/RHN). For detailed values see Supplementary information S2.

|  | A) Means |  |  |  |  |  | B) Differences |  |  |  | *n* |
| --- | --- | --- | --- | --- | --- | --- | --- | --- | --- | --- | --- |
| | overall | | FWB | | RHN | | Δoverall | ΔFWB | ΔRHN | FWB:RHN | |
|  | t1 | t2 | t1 | t2 | t1 | t2 |  |  |  |  |  |
| SO<sub>x</sub> | 21.1 | 8.0 | 17.7 | 8.3 | 24.8 | 7.6 | ↓*** | ↓*** | ↓*** | *** | 40/21/19 |
| NH<sub>y</sub> | 8.7 | 12.5 | 7.7 | 12.2 | 9.7 | 12.7 | ↑*** | ↑*** | ↑*** | *** | 40/21/19 |
| NO<sub>x</sub> | 10.7 | 10.2 | 10.1 | 10.2 | 11.4 | 10.1 | ↓*** | — | ↓*** | *** | 40/21/19 |
| N<sub>total</sub> | 19.4 | 22.6 | 17.8 | 22.5 | 21.1 | 22.8 | ↑*** | ↑*** | ↑*** | *** | 40/21/19 |
| NH<sub>y</sub>:NO<sub>x</sub> | 0.8 | 1.2 | 0.8 | 1.2 | 0.9 | 1.3 | ↑*** | ↑*** | ↑*** | ns | 40/21/19 |

Units: SO<sub>x</sub>: kg S ha<sup>-1</sup> a<sup>-1</sup>; NH<sub>y</sub>, NO<sub>x</sub>, N<sub>tot</sub>: kg N ha<sup>-1</sup> a<sup>-1</sup>

#### Changes in pH and C:N ratio, mean indicator values and vegetation structure

At regional scale, we found an increase in soil pH and a decrease in CN (Table 3). However, the increase in pH was only statistically significant in FWB but not in RHN. ΔpH was negatively correlated with the initial pH (r = −0.59, P < 0.001, *n* = 94). CN decreased in 52 out of 80 plots resulting a decline in both study areas. These changes coincide with general increases in mean R and N indicator values (Table 3). Out of 97 plots, 78 plots showed an increase in mean N values and 68 plots an increase in mean R values. This trend occurred in both study areas, although increases of p/a-based mean R and N were not statistically significant in RHN. There were increases in cover of the shrub and moss layer with a significant difference between the study areas but we did not found no overall change in the herb layer cover at regional scale (Table 3). Trends of shrub and herb layers were different between study areas: In FWB there was an increase in shrub and herb layer cover, whereas the shrub layer remained stable and the herb layer decreased in RHN.

**Table 3.** Soil variables, mean indicator values and structural variables at regional (overall) and local scale (FWB: Fulda-Werra-Bergland, RHN: Rhön Mountains): A) Means for initial survey (t1) and resurvey (t2) and B) Results of exact Wilcox-Tests: One sample tests on differences of variables (Δv_(t2-t1)_) and FWB:RHN: two sample tests on differences between study areas. ↑ or ↓ indicate trends of change, while — indicates no change. Significance levels are indicated by · P < 0.1, * P < 0.05, ** P < 0.01, *** P < 0.001, ns: not significant. *n* = number of plots (overall/FWB/RHN). For detailed values see Supplementary Information S3.

|  | A) Means |  |  |  |  |  | B) Differences |  |  |  | *n* |
| --- | --- | --- | --- | --- | --- | --- | --- | --- | --- | --- | --- |
| | overall | | FWB | | RHN | | Δoverall | ΔFWB | ΔRHN | FWN: RHN | |
|  | t1 | t2 | t1 | t2 | t1 | t2 |  |  |  |  |  |
| Soil parameters |  |  |  |  |  |  |  |  |  |  |  |
| Soil pH | 4.49 | 4.80 | 4.40 | 4.65 | 4.73 | 5.16 | ↑*** | ↑*** | — | * | 94/58/36 |
| Soil C:N ratio | 14.06 | 13.42 | 14.65 | 13.85 | 12.55 | 12.30 | ↓*** | ↓** | ↓*** | ns | 80/51/29 |
| Indicator values |  |  |  |  |  |  |  |  |  |  |  |
| Mean R<br>(cover weighted) | 3.01 | 3.57 | 3.01 | 3.43 | 3.00 | 3.92 | ↑*** | ↑*** | ↑*** | ns | 97/60/37 |
| Mean R<br>(p/a weighted) | 3.52 | 3.73 | 3.48 | 3.59 | 3.64 | 4.07 | ↑*** | ↑*** | ↑. | ns | 97/60/37 |
| Mean N<br>(cover weighted) | 2.71 | 3.05 | 2.70 | 3.00 | 2.72 | 3.20 | ↑*** | ↑*** | ↑** | ns | 97/60/37 |
| Mean N<br>(p/a weighted) | 2.94 | 3.30 | 2.90 | 3.20 | 3.07 | 3.55 | ↑*** | ↑*** | ↑. | ns | 97/60/37 |
| Structural parameters |  |  |  |  |  |  |  |  |  |  |  |
| Cover shrub layer (%) | 0.00 | 2.04 | 0.00 | 2.84 | 0.00 | 0.00 | ↑*** | ↑*** | — | * | 87/60/27 |
| Cover herb layer (%) | 90.35 | 90.59 | 89.31 | 91.31 | 93.00 | 88.75 | — | ↑* | ↓** | *** | 87/60/27 |
| Cover moss layer (%) | 23.01 | 43.65 | 19.98 | 36.94 | 30.75 | 60.75 | ↑*** | ↑*** | ↑*** | ns | 87/60/27 |

#### Changes in total species richness and species groups

We could not detect significant changes in total species richness. This applies also to the richness of vascular plants and bryophytes when analysed separately (results not shown). However, we detected overall qualitative and quantitative changes in almost every species groups (Table 4). Regarding sociological groups, character species and other low-productive grassland species declined in number and abundance, whereas agricultural grassland and fallow species showed an overall increase. The latter groups increased significantly only in FWB. Other low-productive grassland species declined in cover only in RHN but not in FWB. For ecological groups, there was a general increase of nutrient indicators and a significant decrease of acidophytic and basiphytic low-nutrient indicators in both richness and cover. With regard to functional groups, graminoids decreased proportionally in cover but not in richness, whereas forbs increased quantitatively and qualitatively. The Gr:Fo ratio showed a decrease at regional scale but qualitatively only in RHN.

**Table 4.**
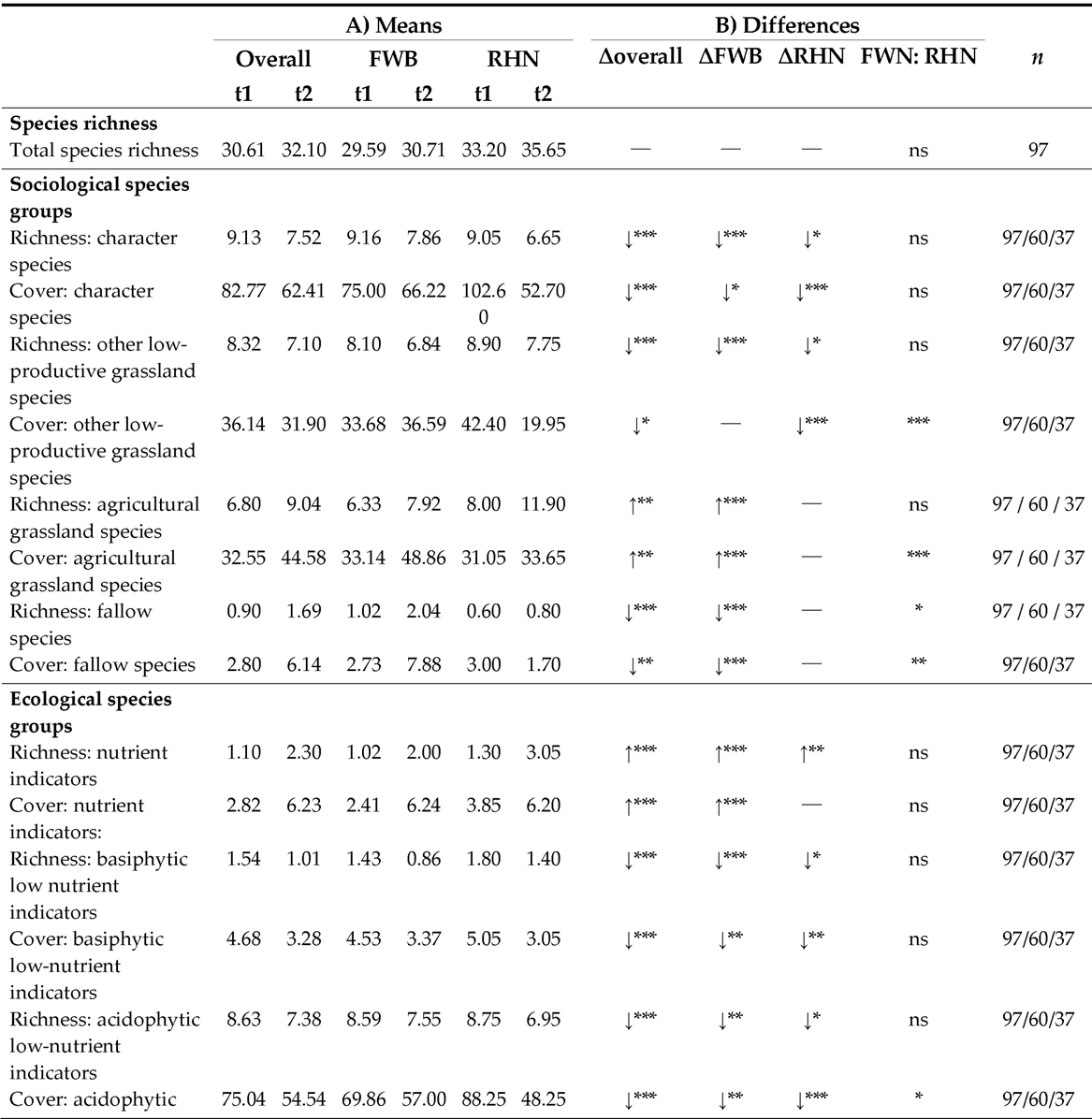
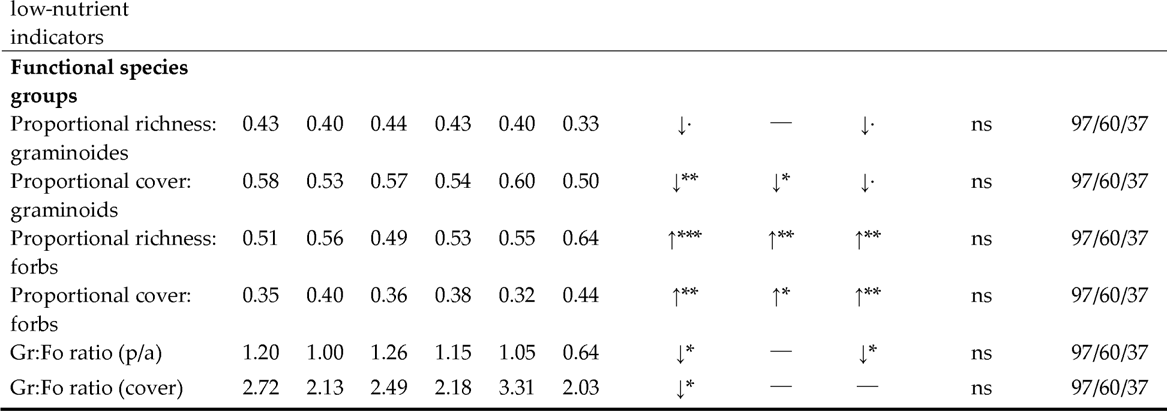
Total species richness, richness and cumulative cover of species groups at regional (overall) and local scale (FWB: Fulda-Werra-Bergland, RHN: Rhön): A) Means for initial survey (t1) and resurvey (t2) and B) Results of exact Wilcox-Tests: One sample tests on differences of variables (Δv_(t2-t1)_) and FWB:RHN: two sample tests on differences between study areas. ↑ or ↓ indicate trends of change, while — indicates no change. Significance levels are indicated by · P < 0.1, * P < 0.05, ** P < 0.01, *** P < 0.001, ns: not significant. *n* = number of plots (overall/FWB/RHN). For detailed values see Supplementary information S3.

#### Changes of individual species: winners and losers

A considerable number of species showed either a significant total increase (14 spp.) or a decrease (19 spp.) in frequency and/or abundance (Supplementary information S4). Declining species belonged mainly to the groups of character species (e.g. *Arnica montana, Danthonia decumbens, Nardus stricta, Calluna vulgaris*) and other low-nutrient indicators (e.g. *Festuca ovina agg., Briza media, Thymus pulegoides, Carex panicea*). Mean R and N indicator values of decreasing species were low with 3.4 for R and 2.5 for N, respectively. Increasing species were notably agricultural grassland species (e.g. *Holcus lanatus, Trifolium pratense, T. repens*) or indifferent species (e.g. *Taraxacum* sect. Ruderalia, *Veronica chamaedrys, Rhytidiadelphus squarrosus*) with higher R and N values (mean 5.3 and 4.9, respectively). Only five species showed different changes between study areas, while four of them occurred only in FWB or RHN, whilst *Campanula rotundifolia*, which occurred in both study areas, showed a significantly stronger decline in FWB than in RHN.

#### Drivers of floristic change: regression models

The results of the regression models indicate which environmental variables had an influence on changes in species composition at the plot level (Table 5, Supplementary information S5). The most important predictor was ΔpH, which influenced most of the dependent variables positively. Only graminoids were negatively related to ΔpH in both richness and cover.

**Table 5.** Multiple linear regression models of environmental change drivers (ΔpH, ΔCN, ΔManagement) and study area on changes in species composition at plot scale. R^2^ and overall significance level of the model. RHN: study area Rhön Mountains in contrast to study area FWB, management change categories: fallow to managed (FM) and continuously managed (MM) in contrast to fallow. ↑ or ↓ indicate trends of regression coefficients. Significance levels are indicated by P < 0.1, * P < 0.05, ** P < 0.01, *** P < 0.001. For detailed values see Supplementary Information S5.

| | $R^2$ | ΔpH | ΔCN | ΔM:FM | ΔM:MM | RHN |
| --- | --- | --- | --- | --- | --- | --- |
| **Mean indicator values** |  |  |  |  |  |  |
| Δmean R value (cover-weighted) | 0.25*** | ↑*** | | | | |
| Δmean R value (p/a) | 0.24*** | ↑*** | | | | |
| Δmean N value (cover-weighted) | | | | | | |
| Δmean N value (p/a) | | | | | | |
| **Species richness** |  |  |  |  |  |  |
| Δtotal richness | 0.20*** | ↑*** | | | | |
| **Sociological species groups** |  |  |  |  |  |  |
| Δrichness character species | | | | | | |
| Δcover character species | 0.06* | | | | | ↓* |
| Δrichness other low-productive grassland species | 0.29*** | ↑*** | | | | ↓*** |
| Δcover other low-productive grassland species | 0.22*** | ↑*** | | | | |
| Δrichness agricultural grassland species | 0.15*** | ↑*** | | | | |
| Δcover agricultural grassland species | 0.13*** | ↑*** | | | | |
| Δrichness fallow species | 0.12** | | | | ↓** | |
| Δcover fallow species | 0.21*** | | | ↓** | ↓*** | |
| **Ecological species groups** |  |  |  |  |  |  |
| Δrichness nutrient indicators | 0.08* | ↑* | | | | |
| Δcover nutrient indicators | 0.08* | ↑* | | | | |
| Δrichness basiphytic low-nutrient indicators | 0.21*** | ↑*** | | | | |
| Δcover basiphytic low-nutrient indicators | 0.20*** | ↑*** | | | | |
| Δrichness acidophytic low-nutrient indicators | | | | | | |
| Δcover acidophytic low-nutrient indicators | 0.08* | | | | | ↓* |
| **Functional species groups** |  |  |  |  |  |  |
| Δrichness graminoids | 0.14** | ↓** | ↓* | | | |
| Δcover graminoids | 0.16** | ↓** | ↓* | | | |
| Δrichness forbs | 0.12** | ↑** | | | | |
| Δcover forbs | 0.15*** | ↑*** | | | | |
| ΔGr:Fo ratio (richness) | | | | | | |
| ΔGr:Fo (cover) | 0.08* | ↑* | | | | |

Contrary to our expectations, we found no effect of ΔpH on mean N indicator values, acidophytic low-nutrient indicators, character species and fallow species. ΔCN had a minor influence with a partial negative effect on graminoids. Ongoing abandonment had a significant positive effect on richness and abundance of fallow species. Study area had no significant influence, except on the cover of character species, on acidophytic low-nutrient indicators and the cover of basiphytic low-nutrient indicators. All of these groups showed significantly higher decreases in RHN than in FWB.

#### Drivers of floristic change: redundancy analysis

Significant effects of ΔpH and ΔM on species composition (cover-weighted and p/a, plot dataset with *n* = 80) have been revealed by the RDA models (with single predictive variables), whereas ΔCN had no effect (Table 6). In most models, study area had a partial effect on species composition, reflecting the different a-priori species compositions of the two study areas. However, study area had no additional effect on p/a species data when ΔM was the constraining variable. An interaction effect with study area was only found for ΔM in cover-weighted species data.

**Table 6.** Results of single variable RDA models of cover-weighted and p/a species data including interaction with study area: A) soil parameters and management change and B) deposition data. Given are the results (P values, significant values with P < 0.05 set in bold) of permutation tests on additional effects of the respected variable (var), effects of study area (SA) and respective interactions.

| **A) Changes in soil parameters and management (plot dataset, *n* = 80)** |  |  |  |  |  |  |
| --- | --- | --- | --- | --- | --- | --- |
|  | *P* (var) | *P* (SA) | *P* (var:SA) | total<br>variance | constrained<br>variance | %-<br>explained |
| **Cover-weighted species data** |  |  |  |  |  |  |
| ΔpH | **0.002** | **0.001** | 0.176 | 139.68 | 10.24 | 7.33 |
| ΔCN | 0.589 | **0.001** | 0.102 | 139.68 | 8.03 | 5.75 |
| Δmanagement | **0.003** | **0.005** | **0.010** | 139.68 | 14.34 | 10.26 |
| **p/a species data** |  |  |  |  |  |  |
| ΔpH | **0.001** | **0.003** | 0.113 | 24.62 | 1.55 | 6.30 |
| ΔCN | 0.918 | **0.004** | 0.692 | 24.62 | 1.01 | 4.09 |
| Δmanagement | **0.006** | 0.123 | 0.288 | 24.62 | 1.89 | 7.67 |
| **B) Changes in deposition (grid cell-pooled dataset, *n* = 40)** |  |  |  |  |  |  |
| **Cover-weighted species data** |  |  |  |  |  |  |
| ΔSO<sub>x</sub> | 0.091 | **0.001** | 0.208 | 139.68 | 8.58 | 6.14 |
| ΔNH<sub>y</sub> | 0.130 | **0.001** | 0.081 | 139.68 | 8.72 | 6.24 |
| ΔNO<sub>x</sub> | 0.152 | **0.001** | **0.002** | 139.68 | 9.04 | 6.47 |
| **p/a species data** |  |  |  |  |  |  |
| ΔSO<sub>x</sub> | **0.004** | **0.036** | 0.300 | 24.62 | 1.23 | 4.98 |
| ΔNH<sub>y</sub> | **0.003** | **0.007** | 0.481 | 24.62 | 1.25 | 5.07 |
| ΔNO<sub>x</sub> | **0.003** | 0.052 | **0.006** | 24.62 | 1.39 | 5.64 |

Based on these results, we calculated the minimal adequate multiple RDA models on cover-weighted and p/a species data which included both ΔpH and FF (as only significant factor level of ΔM) as constraining variables and study area as conditioning variable. In both ordinations, axis 1 reflects species composition shifts connected to changes in soil pH and axis 2 composition shifts connected to management changes, which were however mainly restricted to FWB (Figure 1, Supplementary information S6). Species arrows and correlations with species group changes indicate a positive response of nutrient indicators and/or basiphytes (e.g. *Plantago lanceolata, Rumex acetosa, Thymus pulegioides*) to increasing pH and a negative response of acidophytic species (e.g. *Nardus stricta, Vaccinium myrtillus, acidophytic bryophytes*). Woody species, *Molinia caerulea* and *Galeopsis tetrahit* increased on fallow sites, whereas many low-productive grassland species display a negative trend.

**Figure 1.**
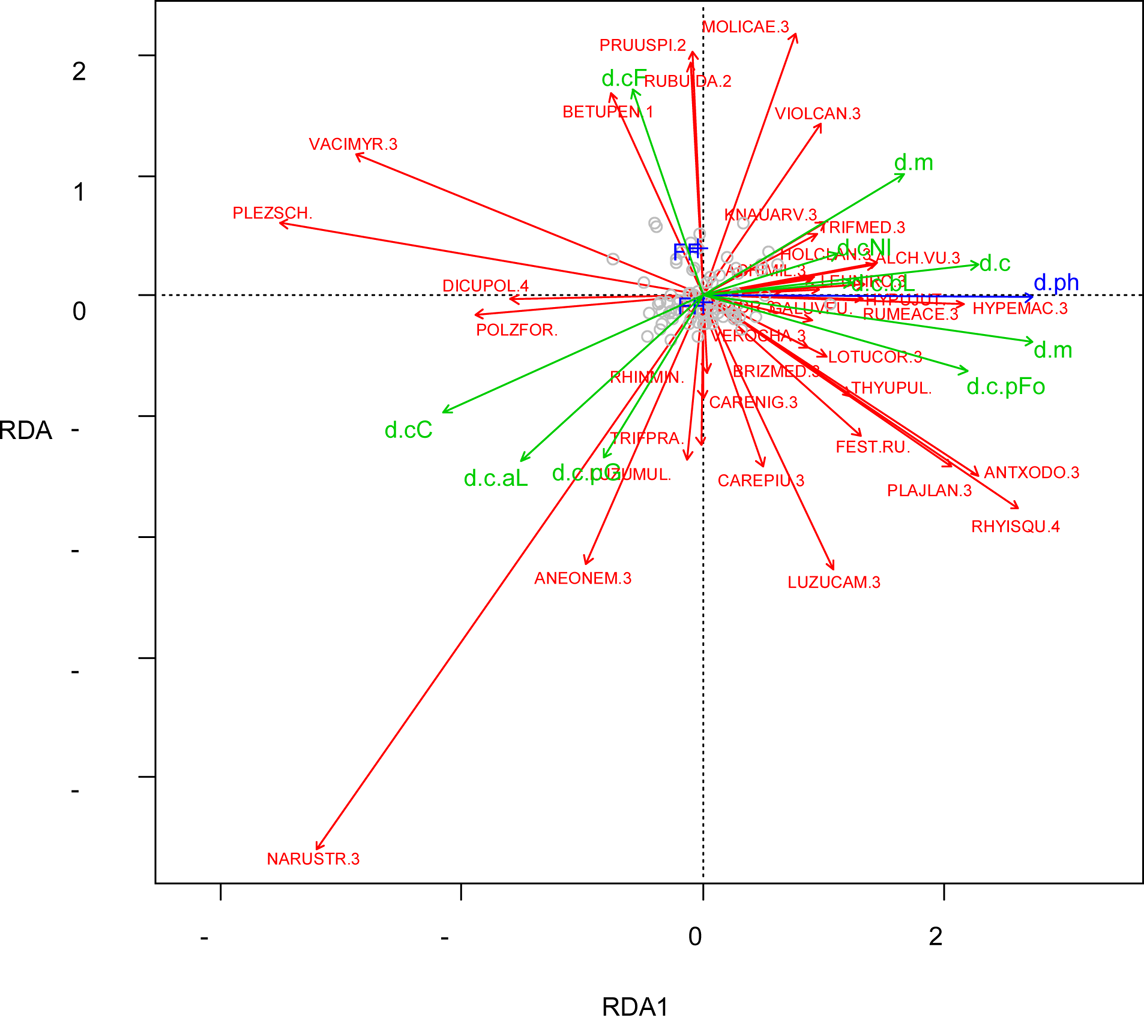
RDA of species differences (cover weighted data) with ΔpH and ΔM (FF vs. FM and MM) as constraining variables. Species data were controlled for study area (conditioning variable). For detailed results see Supplementary information S6. Blue arrows: constraining variables; FF1: fallow at resurvey, FF0: managed at resurvey; ΔpH change in soil pH; Green arrows: supplementary variables (correlations). Abb.: d.c Δ of cover weighted data; mR mean weighted R value; mN mean weighted N value (.pa unweighted); tRi total species richness; C character species; D other low-productive grassland species; G agricultural grassland species; F fallow species; aLI acidophytic low nutrient indicators; bLI basiphytic low nutrient indicators; NI nutrient indicators; pF proportion of forbs; pG proportion of graminoids; .1 tree layer; .2 shrub layer; .3 herb layer; .4 moss layer. Inclusion rules for species: cover-weighted: variance of sqrt(|Δ cover|)-values >0.4 & P-value of multiple linear regression of cover-values on axis 1+2 scores < 0.05, p/a: no. of plots with changes > 4 & P-value of multiple linear regression of p/a values on axis 1+2 scores < 0.05.

*Effects of sulphur and nitrogen deposition on environmental variables and species composition* Regression analyses showed no significant effects of S or N deposition changes (ΔSOx, ΔNHy, ΔNOx) on pooled ΔpH and ΔCN (results not shown). In the RDAs none of the deposition change variables showed additional effects on cover-weighted species change when study area was the first constraining variable (Table 6). In contrast, we found significant effects of ΔNHx and, though less pronounced, ΔSOx for the p/a species change, irrespective of study area. For ΔNOx an interaction effect was detected, indicating that the effect on species composition was restricted to one of the study areas (i.e. RHN).

To combine effects of soil and deposition change (ΔpH and ΔNHy), while taking other factors into account, which influence species composition and the deposition regime, we calculated a multiple RDA model for p/a species data with ΔpH and ΔNHy as constraining variables and ΔM, altitude and study area as conditioning variables. The ordination diagram indicates the floristic pattern reflecting the influence of NHy change (Figure 2). We interpret the first axis as a combined eutrophication effect of ΔpH and ΔNHy, with negative axis 1 scores linked to increasingly occurring nutrient demanding species, a higher proportion of forbs and increasing mean N and R indicator values. Character species and acidophytes react in reverse manner. Axis 2 summarises remaining contrasting effects of both predictor variables, which probably reflect local acidification effects of NHy. This component corresponds to a local spatial windward-leeward pattern of the plots in the Rhön Mountains with more wind-exposed plots on the north-western part (high axis 2 scores) and leeward south-western plots (low axis 2 scores).

**Figure 2.**
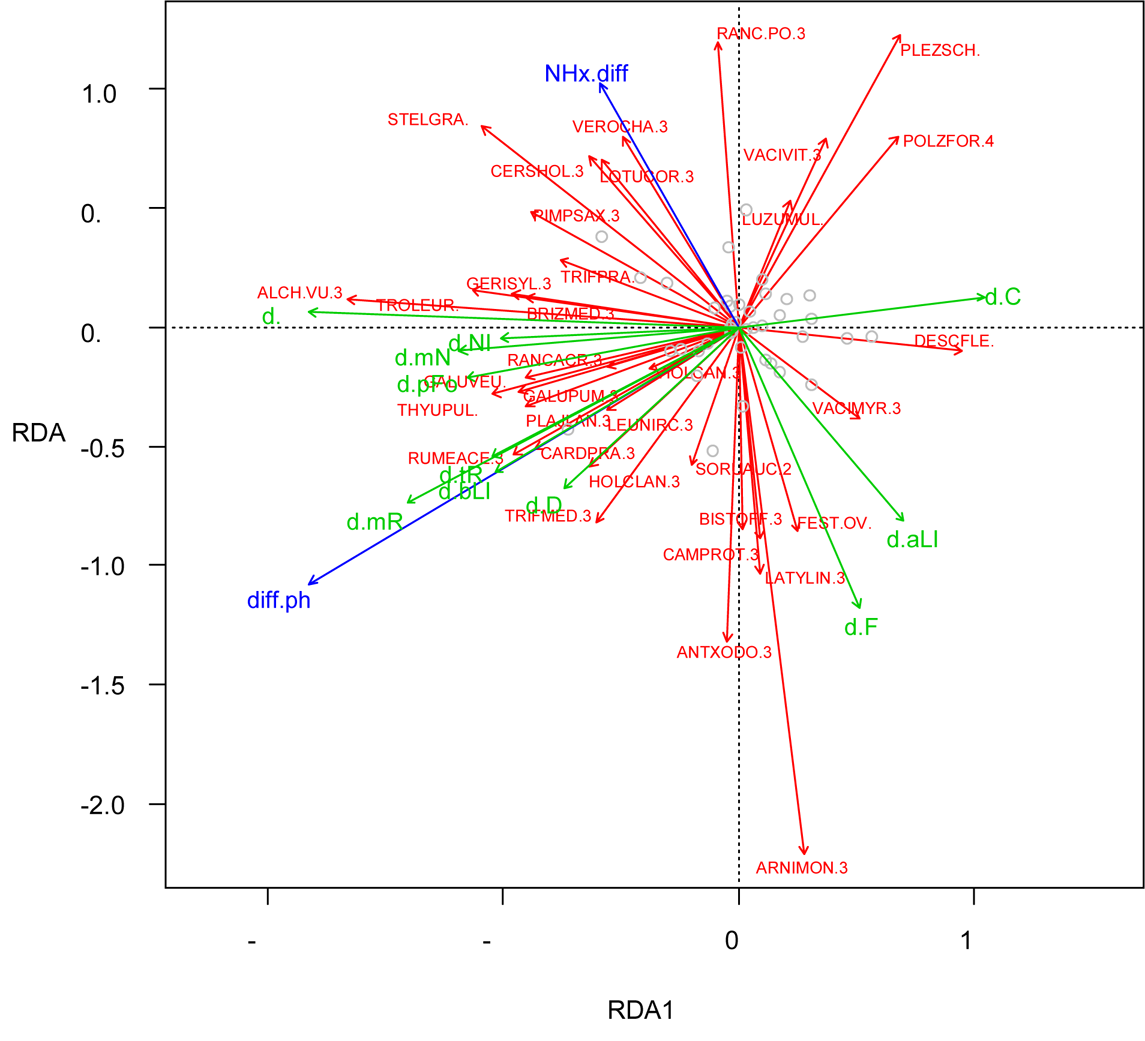
RDA of species differences (p/a-data) with ΔpH and ΔNHy as constraining variables. Species data were controlled for study area, altitude and management (conditioning variables). For detailed results, see Supplementary information S6. Blue arrows: constraining variables; ΔpH change in soil pH, ΔNHy change in reduced N deposition; Green arrows: supplementary variables (correlations); Δ of p/a indicator values and p/a of species groups. Abb.: d. Δ(p/a); mR mean R value; mN mean N value; tRi total species richness; C ri. character species; D ri. other low-productive grassland species; G ri. agricultural grassland species; F ri. fallow species; aLI ri. acidophytic low nutrient indicators; ri. bLI basiphytic low nutrient indicators; ri NI nutrient indicators; pF proportion of forbs; .1 tree layer; .2 shrub layer; .3 herb layer; .4 moss layer. Inclusion rules for species: no. of plots with changes > 4 & P value of multiple linear regression of p/a values on axis 1+2 scores < 0.05.

## Discussion

Our results indicate a strong change towards eutrophication and less acidic conditions of *Nardus* grasslands in parts of the Continental biogeographic region over the past decades. In the vegetation, eutrophication was indicated by increased N indicator values (changes in means, increase in proportion of nutrient indicators, decrease in low-nutrient indicators) and a decrease in CN. The vegetation response to less acidic conditions, indicated by the increase in soil pH, was an increase in mean R indicator values and a decline of acidophytes. Consistently in both study areas, ΔpH demonstrates to be the most important predictor for changes in total species richness and species composition, particularly for the increase of forbs, basiphytes and agricultural grassland species and the decrease of graminoids. Unexpectedly, the significant decline of character species was not related to ΔpH neither at local nor at regional scale. Instead, the decline of character species for plots in FWB was related to a reduced Of-layer (Peppler-Lisbach & Könitz, 2017).

There are several possible causes for eutrophication effects. Direct fertilisation can be ruled out in our study because all plots were either situated within nature reserves and therefore managed without fertilisation or fallow. Therefore, other reasons have to be taken into account. We see three potential drivers for eutrophication: Firstly increased airborne N deposition, secondly reduced SOx deposition with subsequent changes in nutrient availability in soils and thirdly reduced management intensities with processes of auto-eutrophication and insufficient nutrient removal.

The increase of airborne N deposition can be a possible source of eutrophication in *Nardus* grasslands, since total N deposition exceeded the critical loads for this grassland type (10 to 20 kg N ha^-1^ a^-1^), at which the community is expected to lose its stability (Bobbink & Hettelingh, 2011). However, high N deposition rates cannot explain the significant increase in soil pH. On the contrary, increased N deposition would expect further acidification as indicated by declining pH values (Stevens, Duprè, Gaudnik et al., 2011). This is particularly true for NHy depositions, which increased in the study areas during past decades. Expected changes in species composition would be an increase of acid-tolerant species and a decline of species adapted to moderately acid to neutral soil reaction, e.g. many agricultural grassland species (Stevens, Manning et al., 2011). Our findings show, however, the opposite picture: according to the increased soil pH, sites became more favourable for species of agricultural grasslands with higher R and N indicator values, while acidophytic species and low-nutrient indicators declined. The relevant driver for recent changes in soil pH is hence the decrease in SOx deposition rates, particular as there is a strong correlation between atmospheric acid depositions and topsoil pH (Kirk et al., 2010; Stevens et al., 2009). Several other studies report effects of declined SOx deposition rates in Europe since the 1990s on soil properties and more recently on species composition of semi-natural grasslands (e.g. McGovern et al., 2011; Morecroft et al., 2009). Changes in soil pH also result in changes in N availability and soil NH_4_:NO_3_ ratio (Stevens, Manning et al., 2011). Stevens, Duprè, Dorland et al. (2011) predicted that with increasing soil pH, NHy deposition inputs will be progressively converted into nitrate and thus favour N-demanding, acid-intolerant species like those from agricultural grasslands. Although nitrification processes potentially bear the risk of soil acidification, increased pH values indicate that in our case this process is overruled due to decreasing SOx depositions and the soil buffering capacity. Following Rose et al. (2016), Peppler-Lisbach & Könitz (2017) and Mitchell et al. (2018), we suggest therefore that the observed changes of this study are significantly triggered by recovery from acidification due to decreasing SOx depositions. This interpretation would explain the combined pattern of decreasing acidification and eutrophication.

Contrary to our expectations, local patterns of changes in soil parameters were not related to local patterns of N and S deposition. However, we could detect an effect of NHy and SOx deposition change on local patterns of floristic change in the RHN, which may also be the case for differences in floristic changes between FWB and RHN as deposition rates changed differently between study areas and across altitudes. We were not able to separate deposition effects from other drivers differing between study areas. Other studies covering a wider geographical range detected significant common patterns of N deposition and floristic change (Dupré et al., 2010). The reason for the relatively low influence of the local deposition change patterns in our study could be that the overall magnitudes of atmospheric deposition change acts as a master factor that overcompensates local variations. Hence, deposition changes might have triggered the general pattern of floristic changes, but can only poorly explain local differences (Damgaard et al., 2011).

Lastly another factor that contributes to eutrophication are reduced management intensities supporting ‘auto-eutrophication’, i.e. the promotion of tall growing, N-demanding species with an internal nutrient cycling resulting in less nutrient losses through biomass removal (Leuschner & Ellenberg, 2017). The lack of detailed information on temporal management changes between the two surveys and the unbalanced design limit the interpretation of inter-correlations between management history and vegetation change. Nevertheless, we can confirm that on fallow sites fallow indicator species have been favoured and that small herbaceous species have declined accordingly. This pattern has been reported from several studies on secondary succession in *Nardus* grasslands (Schwabe et al., 2019). Interestingly, fallow species did not only increase in fallow plots but were still present in formerly fallow plots, in which management had been re-established prior to the resurvey. This could be due to legacy effects of the fallow period, which are still detectable despite re-introduced management (Arens & Neff, 19970; Hansson & Fogelfors, 2000) or, alternatively, to insufficient suppression of fallow species by current late and low-intensity management (Scheidel & Bruelheide, 2004; Schiefer, 1982). Generally, we have no indication of mediating effects by management to the identified soil and vegetation change, which indicates that the deposition-related drivers and management act rather independently.

The results show widely consistent long-term changes in *Nardus* grasslands in both study areas and confirm local scale findings of Peppler-Lisbach and Könitz (2017). We could hardly detect any contradicting results, except that total herb cover increased in FWB but decreased in RHN. Differences in quantity or significance between study areas relate mainly to changes of some sociological species groups. For example, agricultural grassland species change might not have been significant in RHN because some sociologically indifferent taxa with a significant increase, such as *Taraxacum* Sect. Ruderalia or *Veronica chamaedrys*, are not classifiable to this group. Moreover, an unbalanced distribution of management change classes is mainly responsible for differing significance between study areas. At species level, most species with significant changes displayed the same trend in both study areas and significant differences can largely be explained by the lack of the respective species in the species composition in one of the study areas. The fact that pH did not increase significantly in RHN fits in the general pattern that soils with high initial pH show a less pronounced pH recovery than more acidic soils (Kirk et al., 2010), which were more frequent in FWB. Moreover, the RHN plots included a few plots with organic soils, which tend to show a weak pH recovery due to a lack of weatherable minerals (Kirk et al., 2010).

Generally, a minimum level of inaccuracy in resurveying quasi-permanent plots cannot be entirely excluded and minor differences between study areas could be possibly linked to this methodological issue (Kapfer et al., 2017). However, the ecological consistency of the results suggests that a possible pseudo-turnover plays a minor role in our resurvey study (Ross, Woodin, Hester, Des Thompson, & Birks, 2010; Verheyen et al., 2018).

In light of other European studies, our overall results at regional scale suggest that changes in *Nardus* grasslands follow a general trend across Europe. The detected patterns of eutrophication and recovery from acidification in the Continental biogeographic region are widely consistent with findings from the Atlantic biogeographic region (Mitchell et al., 2018; Stevens, Duprè, Gaudnik et al., 2011). However, beside atmospheric depositions land-use change is still an important trigger. Low management intensities seem to increase species losses and support indirectly eutrophication and the spread of grassland generalists (Rose et al., 2016). Conservation should therefore consider these corresponding drivers in defining their efforts and measures to counteract attested biodiversity losses.

## Conclusion

The findings of this study highlight the risk of eutrophication as a long-term cross-regional threat for species-rich *Nardus* grasslands, which ranges from the Atlantic into the Continental biogeographic region of Europe. Furthermore, they illustrate that both changes in the deposition regime of air pollutants and management have contributed to the detected species turnover. Due to a higher eutrophication pressure, an adapted management becomes an challenging issue. Measures like regular management, earlier mowing dates and moderately increased grazing intensities could become increasingly important to compensate for N depositions and more favourable mineralisation conditions. This underpins the importance of expanded conservation efforts to protect the high biodiversity of *Nardus* grassland under future global change (Stevens et al., 2016). Moreover, the study highlights the relevance of long-term data in combination with resurveys to support our consistent understanding of environmental driver interactions and their impacts on biodiverse semi-natural grasslands.

## Acknowledgements

We would like to thank T. Gauger (Institute of Navigation, University of Stuttgart) for his consult during the analysis of the deposition data and U. v. Borstel (Celle, Germany) for providing his original survey materials and sharing his memories about historic vegetation plots. We also thank D. Meißner, P. Möller and A. Reinhard for help during fieldwork and in processing soils samples. The authors declare to have no conflicts of interest. We thank two anonymous reviewers and the editor for their constructive and helpful comments to improve an earlier version of the manuscript.

## Author contributions

CPL, NK, NS and GR designed the study and conducted the resurvey; CPL and NS analysed the vegetation and environmental data; all authors contributed to the interpretation of the results and discussed them; CPL and NS drafted the manuscript, on which GR and NK gave critical comments and revisions. All authors gave their final approval for publication.

## Supplementary Information

**Supplementary Information S1.** Assignment of species to species groups.

**Supplementary Information S2.** Results of tests on differences in deposition rates.

**Supplementary Information S3.** Results of tests on differences in soil variables, structural variables, Ellenberg indicator values, total species richness, species richness and cover of species groups.

**Supplementary Information S4.** Results of tests on differences in species frequency and cover in the dataset.

**Supplementary Information S5.** Multiple linear regression models of differences in species composition and Ellenberg indicator values on environmental drivers, management change and study area.

**Supplementary Information S6.** Detailed results of RDA analyses of species differences.

## References

1. Arens, R., & Neff, R. (1997). Versuche zum Erhalt von Extensivgrünland: Aus dem wissenschaftlichen Begleitprogramm zum E+E-Vorhaben des Bundesamtes für Naturschutz “Renaturierung des NSG Rotes Moor / Hohe Rhön”. Schriftenreihe Angewandte Landschaftsökologie: H. 13. Bonn-Bad Godesberg.

2. Armstrong, R. H., Grant, S. A., Common, T. G., & Beattie, M. M. (1997). Controlled grazing studies on *Nardus* grassland: Effects of between-tussock sward height and species of grazer on diet selection and intake. Grass and Forage Science, 52(3), 219–231. https://doi.org/10.1111/j.1365-2494.1997.tb02352.x

3. Bobbink, R., & Hettelingh, J.-P. (2011). Review and revision of empirical critical loads and dose-response relationships: Proceedings of an expert workshop, Noordwijkerhout, 23-25 June 2010. Bilthoven: RIVM.

4. Bobbink, R., & Hicks, W. K. (2014). Factors Affecting Nitrogen Deposition Impacts on Biodiversity: An Overview. In M. A. Sutton, K. E. Mason, L. J. Sheppard, H. Sverdrup, R. Haeuber, & W. K. Hicks (Eds.), Nitrogen Deposition, Critical Loads and Biodiversity (pp. 127–138). Dordrecht: Springer Netherlands. https://doi.org/10.1007/978-94-007-7939-6_14

5. Bobbink, R., Hicks, K., Galloway, J., Spranger, T., Alkemade, R., Ashmore, M. R., Vries, W. de (2010). Global assessment of nitrogen deposition effects on terrestrial plant diversity: A synthesis. Ecological Applications, 20(1), 30–59. https://doi.org/10.1890/08-1140.1

6. Bobbink, R., Hornung, M., & Roelofs, J. G. M. (1998). The effects of air-borne nitrogen pollutants on species diversity in natural and semi-natural European vegetation. Journal of Ecology, 86(5), 717–738. https://doi.org/10.1046/j.1365-2745.1998.8650717.x

7. Bohn, U., Korneck, D., & Meisel, K. (1996). Vegetationskarte der Bundesrepublik Deutschland 1:200000: Blatt CC 5518 Fulda (2nd ed.). Schriftenreihe für Vegetationskunde: Vol. 15. Bonn-Bad Godesberg: BfN.

8. Borstel, U.-O. von. (1974). Untersuchungen zur Vegetationsentwicklung auf ökologisch verschiedenen Grünland-und Ackerbrachen hessischer Mittelgebirge (Westerwald, Rhön, Vogelsberg) (Dissertation). Justus-Liebig-Universität, Gießen.

9. Ceulemans, T., Merckx, R., Hens, M., & Honnay, O. (2013). Plant species loss from European semi-natural grasslands following nutrient enrichment - is it nitrogen or is it phosphorus? Global Ecology and Biogeography, 22(1), 73–82. https://doi.org/10.1111/j.1466-8238.2012.00771.x

10. Damgaard, C., Jensen, L., Frohn, L. M., Borchsenius, F., Nielsen, K. E., Ejrnæs, R., & Stevens, C. J. [Carly J.] (2011). The effect of nitrogen deposition on the species richness of acid grasslands in Denmark: A comparison with a study performed on a European scale. Environmental Pollution, 159(7), 1778–1782. https://doi.org/10.1016/j.envpol.2011.04.003

11. De Graaf, M. C. C., Bobbink, R., Roelofs, J. G. M., & Verbeek, P. J. M. (1998). Differential effects of ammonium and nitrate on three heathland species. Plant Ecology, 135(2), 185– 196. https://doi.org/10.1023/A:1009717613380

12. Dengler, J. [Juergen], Janisova, M., Toeroek, P., & Wellstein, C. (2014). Biodiversity of Palaearctic grasslands: A synthesis. Agriculture Ecosystems & Environment, 182, 1–14. https://doi.org/10.1016/j.agee.2013.12.015

13. Dentener, F., Drevet, J., Lamarque, J. F., Bey, I., Eickhout, B., Fiore, A. M., Wild, O. (2006). Nitrogen and sulfur deposition on regional and global scales: A multimodel evaluation. Global Biogeochemical Cycles, 20(4), n/a-n/a. https://doi.org/10.1029/2005GB002672

14. Dupré, C., Stevens, C. J. [Carly J.], Ranke, T., Bleeker, A. [Albert], Peppler-Lisbach, C., Gowing, D. J. G. [David J. G.], Diekmann, M. (2010). Changes in species richness and composition in European acidic grasslands over the past 70 years: The contribution of cumulative atmospheric nitrogen deposition. Global Change Biology, 16(1), 344–357. https://doi.org/10.1111/j.1365-2486.2009.01982.x

15. Ellenberg, H. (1992). Zeigerwerte von Pflanzen in Mitteleuropa (2nd ed.). Scripta geobotanica. Göttingen: Goltze. European Council. (1992). Council Directive 92/43/EEC of 21 May 1992 on the conservation of natural habitats and of wild fauna and flora.

16. Fennema, F. (1991). SO2 and NH3 deposition as possible causes for the extinction of Arnica montana L. Water, Air, & Soil Pollution, 62(3), 325–336. https://doi.org/10.1007/BF00480264

17. Field, C. D. [Chris D.], Dise, N. B., Payne, R. J., Britton, A. J., Emmett, B. A. [Bridget A.], Helliwell, R. C., Caporn, S. J. M. (2014). The Role of Nitrogen Deposition in Widespread Plant Community Change Across Semi-natural Habitats. Ecosystems, 17(5), 864–877. https://doi.org/10.1007/s10021-014-9765-5

18. Galvánek, D., & Janák, M. (2008). Management of Natura 2000 habitats *Species-rich Nardus grasslands 6230: Technical Report 2008 14/24: European Commission.

19. Gauger, T. (2010). Erfassung, Prognose und Bewertung von Stoffeinträgen und ihren Wirkungen in Deutschland MAPESI-Projekt (Modelling of Air Pollutants and EcoSystem Impact): Kartierung von Deposition Loads 2004 bis 2007, Anhang XI: Textteil und Ergebnis-Statistik: Abschlussbericht im Auftrag des Umweltbundesamtes BMU/UBA 3707 64 200, UBA-Texte 42/2011.

20. Gauger, T., Anshelm, F., Schuster, H., Erisman, J. W. [J. W.], Vermeulen, A. T., Draaijers, G.P.J., Nagel, H.-D. [H.-D.]. (2002). Mapping of ecosystem specific long-term trends in deposition loads and concentrations of air pollutants in Germany and their comparison with Critical Loads and Critical Levels. Berlin: Final Report on behalf of Federal Environmental Agency (Umweltbundesamt), BMU/UBA FE-No 299 42 210.

21. Gauger, T., Haenel, H. D., Rösemann, C., Dämmgen, U., Bleeker, A. [A.], Erisman, J. W. [J. W.], Duyzer, J. H. (2008). National Implementation of the UNECE Convention on Long-range Transboundary Air Pollution (Effects) / Nationale Umsetzung UNECE-Luftreinhaltekonvention (Wirkungen): Part 1: Deposition Loads: Methods, modelling and mapping results, trends.: BMU/UBA 204 63 252. UBA-Texte 38/08 (1). ISSN 1862-4804.

22. Gauger, T., Kölbe, R., & Anshelm, F. (2000). Kritische Luftschadstoff-Konzentrationen und Eintragsraten sowie ihre Überschreitung für Wald-und Agrarökosysteme sowie naturnahe waldfreie Ökosysteme. Teil 1: Deposition Loads. Teil 2: Critical Levels. Universität Stuttgart: Forschungsvorhaben im Auftrag des BMU/UBA, FE-Nr. 297 85 079, Institut für Navigation.

23. Gauger, T., Nagel, H.-D. [Hans-Dieter], Schlutow, A., & Scheuchner, T. (2013). Erstellung einer methodenkonsistenten Zeitreihe von Stoffeinträgen und Ihren Wirkungen in Deutschland: Teil 2 Abschlussbericht. Dessau-Roßlau.

24. Hansson, M., & Fogelfors, H. (2000). Management of a semi-natural grassland; results from a 15-year-old experiment in southern Sweden. Journal of Vegetation Science, 11(1), 31–38. https://doi.org/10.2307/3236772

25. Hejcman, M., Hejcmanova, P., Pavlu, V., & Benes, J. (2013). Origin and history of grasslands in Central Europe - a review. Grass and Forage Science, 68(3), 345–363. https://doi.org/10.1111/gfs.12066

26. Helsen, K., Ceulemans, T., Stevens, C. J. [Carly J.], & Honnay, O. (2014). Increasing Soil Nutrient Loads of European Semi-natural Grasslands Strongly Alter Plant Functional Diversity Independently of Species Loss. Ecosystems, 17(1), 169–181. https://doi.org/10.1007/s10021-013-9714-8

27. Henrys, P. A. [P. A.], Stevens, C. J. [C. J.], Smart, S. M. [S. M.], Maskell, L. C. [L. C.], Walker, K. J., Preston, C. D., Emmett, B. A. [B. A.] (2011). Impacts of nitrogen deposition on vascular plants in Britain: An analysis of two national observation networks. Biogeosciences, 8(12), 3501–3518. https://doi.org/10.5194/bg-8-3501-2011

28. Hothorn, T., & Hornik, K. (2017). exactRankTests.

29. Humbert, J.-Y., Dwyer, J. M., Andrey, A., & Arlettaz, R. (2016). Impacts of nitrogen addition on plant biodiversity in mountain grasslands depend on dose, application duration and climate: A systematic review. Global Change Biology, 22(1), 110–120. https://doi.org/10.1111/gcb.12986

30. Jansen, F., & Dengler, J. [J.] (2008). GermanSL – Eine universelle taxonomische Referenzliste für Vegetationsdatenbanken in Deutschland. Tuexenia, 28, 239–253.

31. Kapfer, J., Hédl, R., Jurasinski, G., Kopecký, M., Schei, F. H., & Grytnes, J.-A. [John-Arvid] (2017). Resurveying historical vegetation data opportunities and challenges. Applied Vegetation Science, 20(2), 164–171. https://doi.org/10.1111/avsc.12269

32. Kirk, G. J.D., Bellamy, P. H., & Murray Lark, R. (2010). Changes in soil pH across England and Wales in response to decreased acid deposition. Global Change Biology, 437, no-no. https://doi.org/10.1111/j.1365-2486.2009.02135.x

33. Kleijn, D., Bekker, R. M., Bobbink, R., de Graaf, M. C. C., & Roelofs, J. G. M. (2008). In search for key biogeochemical factors affecting plant species persistence in heathland and acidic grasslands: a comparison of common and rare species. Journal of Applied Ecology, 45(2), 680–687. https://doi.org/10.1111/j.1365-2664.2007.01444.x

34. Klink, H.-J. (1969). Die naturräumlichen Einheiten auf Blatt 112 Kassel. Geographische Landesaufnahme 1: 200.000. Bundesforschungsanstalt für Landeskunde und Raumordnung. Bonn, Bad Godesberg.

35. Leuschner, C., & Ellenberg, H. (2017). Ecology of Central European Non-Forest Vegetation: Coastal to Alpine, Natural to Man-Made Habitats, Vol II (L. Sutcliffe, Trans.) (Vol II). Cham: Springer International.

36. Mariotte, P., Buttler, A., Kohler, F., Gilgen, A. K., & Spiegelberger, T. (2013). How do subordinate and dominant species in semi-natural mountain grasslands relate to productivity and land-use change? Basic and Applied Ecology, 14(3), 217–224. https://doi.org/10.1016/j.baae.2013.02.003

37. Maskell, L. C. [Lindsay C.], Smart, S. M. [Simon M.], Bullock, J. M., Thompson, K., & Stevens, C. J. [Carly J.] (2010). Nitrogen deposition causes widespread loss of species richness in British habitats. Global Change Biology, 16(2), 671–679. https://doi.org/10.1111/j.1365-2486.2009.02022.x

38. McGovern, S., Evans, C. D., Dennis, P., Walmsley, C., & McDonald, M. A. (2011). Identifying drivers of species compositional change in a semi-natural upland grassland over a 40-year period. Journal of Vegetation Science, 22(2), 346–356. https://doi.org/10.1111/j.1654-1103.2011.01256.x

39. Mitchell, R. J., Hewison, R. L., Fielding, D. A., Fisher, J. M., Gilbert, D. J., Hurskainen, S., Riach, D. (2018). Decline in atmospheric sulphur deposition and changes in climate are the major drivers of long-term change in grassland plant communities in Scotland. Environmental Pollution, 235, 956–964. https://doi.org/10.1016/j.envpol.2017.12.086

40. Morecroft, M. D. [M. D.], Bealey, C. E., Beaumont, D. A. [D. A.], Benham, S. [S.], Brooks, D. R., Burt, T. P., Watson, H. [H.] (2009). The UK Environmental Change Network: Emerging trends in the composition of plant and animal communities and the physical environment. Biological Conservation, 142(12), 2814–2832. https://doi.org/10.1016/j.biocon.2009.07.004

41. Oksanen, J., Blanchet, F. G., Kindt, R., Legendre, P., Minchin, P. R., O’Hara, R. B., Wagner, H. (2015). Vegan.

42. Pakeman, R. J., Alexander, J., Brooker, R., Cummins, R., Fielding, D. A., Gore, S., Lewis, R. (2016). Long-term impacts of nitrogen deposition on coastal plant communities. Environmental Pollution, 212, 337–347. https://doi.org/10.1016/j.envpol.2016.01.084

43. Payne, R. J., Dise, N. B., Field, C. D., Dore, A. J., Caporn, S. J. M., & Stevens, C. J. [Carly J.] (2017). Nitrogen deposition and plant biodiversity: Past, present, and future. Frontiers in Ecology and the Environment, 15(8), 431–436. https://doi.org/10.1002/fee.1528

44. Peppler, C. (1992). Die Borstgrasrasen (Nardetalia) Westdeutschlands. Dissertationes botanicae 193. Dissertationes botanicae: Vol 193. Göttingen, Berlin: Cramer.

45. Peppler-Lisbach, C., & Könitz, N. (2017). Vegetation changes in *Nardus* grasslands of the Werra-Meissner region (Hesse, Lower Saxony, Central Germany) after 25 years. Tuexenia, 37, 201–228. https://doi.org/10.14471/2017.37.001

46. Peppler-Lisbach, C., & Petersen, J. (2001). Synopsis der Pflanzengesellschaften Deutschlands: Calluno-Ulicetea (G3), Teil 1: Nardetalia strictae Borstgrasrasen. Synopsis der Pflanzengesellschaften Deutschlands: H 8. Göttingen.

47. Phoenix, G. K., Emmett, B. A. [Bridget A.], Britton, A. J., Caporn, S. J. M., Dise, N. B., Helliwell, R. C., Power, S. A. (2012). Impacts of atmospheric nitrogen deposition: Responses of multiple plant and soil parameters across contrasting ecosystems in long-term field experiments. Global Change Biology, 18(4), 1197–1215. https://doi.org/10.1111/j.1365-2486.2011.02590.x

48. R Core Team. (2017). ’R’. Vienna, Austria. URL http://www.R-project.org/.

49. Rose, R., Monteith, D. T. [Don T.], Henrys, P. A. [Peter A.], Smart, S. M. [Simon M.], Wood, C., Morecroft, M. [Mike], Watson, H. [Helen] (2016). Evidence for increases in vegetation species richness across UK Environmental Change Network sites linked to changes in air pollution and weather patterns. Ecological Indicators, 68, 52–62. https://doi.org/10.1016/j.ecolind.2016.01.005

50. Ross, L. C., Woodin, S. J., Hester, A., Des Thompson, B. A., & Birks, H. J. B. (2010). How important is plot relocation accuracy when interpreting re-visitation studies of vegetation change? Plant Ecology & Diversity, 3(1), 1–8. https://doi.org/10.1080/17550871003706233

51. Ross, L. C., Woodin, S. J., Hester, A. J., Des Thompson, B. A., Birks, H. J. B., & Wildi, O. (2012). Biotic homogenization of upland vegetation: Patterns and drivers at multiple spatial scales over five decades. Journal of Vegetation Science, 23(4), 755–770. https://doi.org/10.1111/j.1654-1103.2012.01390.x

52. Roy, P.-O., Azevedo, L. B., Margni, M., van Zelm, R., Deschenes, L., & Huijbregts, M. A. J. (2014). Characterization factors for terrestrial acidification at the global scale: A systematic analysis of spatial variability and uncertainty. The Science of the Total Environment, 500, 270–276. https://doi.org/10.1016/j.scitotenv.2014.08.099

53. Sala, O. E., Chapin, F. S., Armesto, J. J., Berlow, E., Bloomfield, J., Dirzo, R., Wall, D. H. (2000). Biodiversity - Global biodiversity scenarios for the year 2100. Science, 287(5459), 1770–1774. https://doi.org/10.1126/science.287.5459.1770

54. Scheidel, U., & Bruelheide, H. (2004). Versuche zur Beweidung von Bergwiesen im Harz. Hercynia N.F., 37, 87–101.

55. Schiefer, J. (1982). Einfluß der Streuzersetzung auf die Vegetationsentwicklung brachliegender Rasengesellschaften. Tuexenia, 2, 209–218.

56. Schwabe, A., Tischew, S., Bergmeier, E., Garve, E., Härdtle, W., Heinken, T., Dierschke, H. (2019). Plant community of the year 2020: Mat grassland (*Nardus* stricta grassland). Tuexenia, 39, 287–308. https://doi.org/10.14471/2019.39.017

57. Southon, G. E., Field, C., Caporn, S. J. M., Britton, A. J., & Power, S. A. (2013). Nitrogen Deposition Reduces Plant Diversity and Alters Ecosystem Functioning: Field-Scale Evidence from a Nationwide Survey of UK Heathlands. Plos One, 8(4). https://doi.org/10.1371/journal.pone.0059031

58. Stevens, C. J. [Carly J.], Dise, N. B., & Gowing, D. J. G. [David J. G.] (2009). Regional trends in soil acidification and exchangeable metal concentrations in relation to acid deposition rates. Environmental Pollution, 157(1), 313–319. https://doi.org/10.1016/j.envpol.2008.06.033

59. Stevens, C. J. [Carly J.], Dise, N. B., Gowing, D. J. G. [David J. G.], & Mountford, J. O. (2006). Loss of forb diversity in relation to nitrogen deposition in the UK: Regional trends and potential controls. Global Change Biology, 12(10), 1823–1833. https://doi.org/10.1111/j.1365-2486.2006.01217.x

60. Stevens, C. J. [Carly J.], Dise, N. B., Mountford, J. O., & Gowing, D. J. G. [David J. G.] (2004). Impact of Nitrogen Deposition on the Species Richness of Grasslands. Science, 303(5665), 1876–1879. https://doi.org/10.1126/science.1094678

61. Stevens, C. J. [Carly J.], Duprè, C., Dorland, E., Gaudnik, C., Gowing, D. J. G. [David J. G.], Bleeker, A. [Albert], Dise, N. B. (2010). Nitrogen deposition threatens species richness of grasslands across Europe. Environmental Pollution, 158(9), 2940–2945. https://doi.org/10.1016/j.envpol.2010.06.006

62. Stevens, C. J. [Carly J.], Duprè, C., Dorland, E., Gaudnik, C., Gowing, D. J. G. [David J. G.], Bleeker, A. [Albert], Dise, N. B. (2011). The impact of nitrogen deposition on acid grasslands in the Atlantic region of Europe. Environmental Pollution, 159(10), 2243–2250. https://doi.org/10.1016/j.envpol.2010.11.026

63. Stevens, C. J. [Carly J.], Duprè, C., Gaudnik, C., Dorland, E., Dise, N. B., Gowing, D. J. G. [David J. G.], Diekmann, M. (2011). Changes in species composition of European acid grasslands observed along a gradient of nitrogen deposition. Journal of Vegetation Science, 22(2), 207–215. https://doi.org/10.1111/j.1654-1103.2010.01254.x

64. Stevens, C. J. [Carly J.], Manning, P., van den Berg Leon, J L, de Graaf, M. C. C., Wamelink, G. W. W., Boxman, A. W., Dorland, E. (2011). Ecosystem responses to reduced and oxidised nitrogen inputs in European terrestrial habitats. Environmental Pollution, 159(3), 665–676. https://doi.org/10.1016/j.envpol.2010.12.008

65. Stevens, C. J. [Carly J.], Payne, R. J., Kimberley, A., & Smart, S. M. [Simon M.] (2016). How will the semi-natural vegetation of the UK have changed by 2030 given likely changes in nitrogen deposition? Environmental Pollution, 208, 879–889. https://doi.org/10.1016/j.envpol.2015.09.013

66. Stevens, C. J. [Carly J.], Thompson, K., Grime, J. P., Long, C. J., & Gowing, D. J. G. [David J. G.] (2010). Contribution of acidification and eutrophication to declines in species richness of calcifuge grasslands along a gradient of atmospheric nitrogen deposition. Functional Ecology, 24(2), 478–484. https://doi.org/10.1111/j.1365-2435.2009.01663.x

67. Teufel, J., Gauger, T., & Braun, B. (1994). Einfluß von Immisionen und Depositionen von Luftverunreinigungen auf Borstgrasrasen in der Bundesrepublik Deutschland (Abschlußbericht FE-Vorhaben Nr. 108 02 101). Stuttgart.

68. Verheyen, K., Bažány, M., Chećko, E., Chudomelová, M., Closset-Kopp, D., Czortek, P., Baeten, L. (2018). Observer and relocation errors matter in resurveys of historical vegetation plots. Journal of Vegetation Science, 29(5), 812–823. https://doi.org/10.1111/jvs.12673

