## Supplementary Information S1 for "Long-term vegetation changes in species-rich *Nardus* grasslands of Central Germany indicate eutrophication, recovery from acidification and management change as main drivers"

**Appendix S1.** Assignment of species to species groups

**Legend**

**Sociological species groups**

C – character species (*Nardetalia* specialists in grassland habitats) (Peppler-Lisbach & Petersen 2001)

D – other low-productive grassland species (species of anthropo-zoogenic heathlands/grasslands and an Ellenberg N indicator value of < 4 (Ellenberg 1992)

G – agricultutal grassland species (species of anthropo-zoogenic heathlands/grasslands and Ellenberg N ≥ 4) (Ellenberg 1992)

F – fallow species (all tree and shrub species and herbaceous species of forests, forest clearings and fringes) (Ellenberg 1992)

All other species: I = indifferent

**Functional species groups**

Fo – Forbs (herbaceous, non-graminoid species)

Gr – Graminoides (Poaceace, Cyperaceae and Juncaceae)

All other species: I = other species (e.g. trees, shrubs, lichens, mosses)

**Ecological species groups**

NI – nutrient indicators (Ellenberg N > 5)

bLI – basiphytic low-nutrient indicators (Ellenberg R >5 & N < 4)

aLI – acidophytic low-nutrient indicators (Ellenberg R < 4 & N < 4

All other species: I = indifferent

| **Full species name** | **Species code** | **Sociological species group** | **Functional species group** | **Ecological species group** |
| --- | --- | --- | --- | --- |
| Acer pseudoplatanus | ACERPSE | F | I | NI |
| Achillea millefolium | ACHIMIL | G | Fo | I |
| Achillea ptarmica | ACHIPTA | G | Fo | I |
| Agrostis canina | AGRTCAN | D | Gr | aLI |
| Agrostis capillaris | AGRTCAP | G | Gr | I |
| Agrostis gigantea | AGRTGIG | G | Gr | NI |
| Agrostis stolonifera | AGRTSTO | G | Gr | I |
| Ajuga reptans | AJUGREP | I | Fo | NI |
| Alchemilla glaucescens | ALCH.VU | G | Fo | I |
| Alchemilla vulgaris agg. | ALCHGLU | D | Fo | I |
| Alopecurus pratensis | ALOPPRA | G | Gr | NI |
| Anemone nemorosa | ANEONEM | I | Fo | I |
| Angelica sylvestris | ANGESYL | G | Fo | I |
| Antennaria dioica | ANTEDIO | D | Fo | aLI |
| Anthoxanthum odoratum | ANTXODO | I | Gr | I |
| Anthriscus sylvestris | ANTUSYL | G | Fo | NI |
| Arnica montana | ARNIMON | C | Fo | aLI |
| Arrhenatherum elatius | ARRHELA | G | Gr | NI |
| Betonica officinalis | BETOOFF | G | Fo | I |
| Betula pendula | BETUPEN | F | I | I |
| Betula pubescens | BETUPUB | F | I | aLI |
| Bistorta officinalis | BISTOFF | G | Fo | I |
| Briza media | BRIZMED | D | Gr | I |
| Calamagrostis epigeios | CALAEPI | F | Gr | NI |
| Calluna vulgaris | CALQVUL | C | S | aLI |
| Caltha palustris | CALWPAL | G | Fo | NI |
| Campanula rotundifolia | CAMPROT | D | Fo | I |
| Cardamine pratensis | CARDPRA | I | Fo | I |
| Cares pilulifera | CAREPIU | C | Gr | aLI |
| Carex canescens | CARECAN | D | Gr | I |
| Carex caryophyllea | CARECAR | D | Gr | I |
| Carex demissa | CAREDEM | D | Gr | I |
| Carex echinata | CAREECH | D | Gr | aLI |
| Carex nigra | CARENIG | D | Gr | aLI |
| Carex ovalis | CAREOVA | C | Gr | aLI |
| Carex pallescens | CAREPAL | C | Gr | I |
| Carex panicea | CAREPAN | D | Gr | I |
| Carex pulicaris | CAREPUL | D | Gr | I |
| Carlina acaulis | CARLA.S | D | Fo | I |
| Carlina vulgaris | CARLVUL | D | Fo | bLI |
| Carpinus betulus | CARIBET | F | I | I |
| Centaurea jacea | CENTJAC | D | Fo | I |
| Centaurea montana | CENTMON | I | Fo | NI |
| Centaurea nigra ssp. nemoralis | CENTN.N | D | Fo | bLI |
| Cerastium arvense | CERSARV | I | Fo | I |
| Cerastium holosteoides | CERSHOL | G | Fo | I |
| Chaerophyllum hirsutum | CHARH.H | I | Fo | NI |
| Cirsium acaule | CIRSACA | D | Fo | bLI |
| Cirsium palustre | CIRSPAL | G | Fo | I |
| Cirsium vulgare | CIRSVUL | I | Fo | NI |
| Colchicum autumnale | COLHAUT | G | Fo | I |
| Convallaria majalis | CONVMAJ | I | Fo | I |
| Corylus avellana | CORDAVE | F | I | I |
| Crataegus laevigata | CRATLAE | F | I | I |
| Crataegus monogyna | CRATMON | F | I | I |
| Crepis mollis | CREPMOL | G | Fo | I |
| Crepis paludosa | CREPPAL | G | Fo | NI |
| Cynosurus cristatus | CYNSCRI | G | Gr | I |
| Cytisus scoparius | CYTSSCO | F | I | I |
| Dactylis glomerata | DACYGLO | G | Gr | NI |
| Dactylorhiza fuchsii | DACLFUC | D | Fo | I |
| Dactylorhiza majalis | DACLMAJ | G | Fo | bLI |
| Danthonia decumbens | DANTDEC | C | Gr | aLI |
| Deschampsia cespitosa | DESCCES | I | Gr | I |
| Deschampsia flexuosa | DESCFLE | C | Gr | aLI |
| Dianthus superbus | DIANS.S | D | Fo | bLI |
| Digitalis purpurea | DIGIPUR | F | Fo | NI |
| Dryopteris carthusiana | DRYPCAR | F | Fo | I |
| Dryopteris filix-mas | DRYPFIL | F | Fo | NI |
| Epilobium angustifolium | EPIOANG | F | Fo | NI |
| Epilobium palustre | EPIOPAL | D | Fo | aLI |
| Equisetum arvense | EQUTARV | I | Fo | I |
| Equisetum sylvaticum | EQUTSYL | F | Fo | I |
| Eriophorum angustifolium | ERIOANG | D | Gr | I |
| Eriophorum vaginatum | ERIOVAG | D | Gr | aLI |
| Euphorbia cyparissias | EUPHCYP | D | Fo | I |
| Euphrasia officinalis ssp. rostkoviana | EUPRO.R | G | Fo | I |
| Euphrasia stricta | EUPRSTR | D | Fo | I |
| Festuca ovina agg. | FEST.OV | D | Gr | I |
| Festuca pratensis | FESTPRA | G | Gr | NI |
| Festuca rubra agg. | FEST.RU | G | Gr | I |
| Filipendula ulmaria | FILIULM | G | Fo | I |
| Frangula alnus | FRANALN | F | I | I |
| Fraxinus excelsior | FRAXEXC | F | I | NI |
| Galeopsis tetrahit | GALOTET | F | Fo | NI |
| Galium album | GALUALB | G | Fo | I |
| Galium aparine | GALUAPA | I | Fo | NI |
| Galium boreale | GALUBOR | D | Fo | bLI |
| Galium palustre ssp. palustre | GALUP.P | I | Fo | I |
| Galium pumilum | GALUPUM | D | Fo | I |
| Galium saxatile | GALUSAX | C | Fo | aLI |
| Galium uliginosum | GALUULI | D | Fo | I |
| Galium verum | GALUVEU | D | Fo | bLI |
| Genista tinctoria | GENITIN | D | L | bLI |
| Geranium sylvaticum | GERISYL | G | Fo | NI |
| Geum rivale | GEUMRIV | G | Fo | I |
| Helianthemum obscurum | HELIN.O | D | Fo | bLI |
| Helictotrichon pratense | HELTPRA | D | Gr | I |
| Helictotrichon pubescens | HELTPUB | G | Gr | I |
| Heracleum sphondylium | HERASPH | G | Fo | NI |
| Hieracium lachenalii | HIERLAC | D | Fo | I |
| Hieracium lactucella | HIERLAT | C | Fo | I |
| Hieracium laevigatum | HIERLAV | D | Fo | aLI |
| Hieracium murorum | HIERMUR | I | Fo | I |
| Hieracium pilosella | HIERPIO | D | Fo | I |
| Holcus lanatus | HOLCLAN | G | Gr | I |
| Holcus mollis | HOLCMOL | F | Gr | aLI |
| Hypericum humifusum | HYPEHUM | D | Fo | I |
| Hypericum maculatum | HYPCMAC | D | Fo | bLI |
| Hypericum perforatum | HYPEPEF | D | Fo | I |
| Hypnum jutlandicum | HYPUJUT | C | I | I |
| Hypochaeris maculata | HYPEMAC | D | Fo | aLI |
| Hypochaeris radicata | HYPCRAD | D | Fo | I |
| Juncus acutiflorus | JUNUACU | D | Gr | I |
| Juncus conglomeratus | JUNUCON | D | Gr | I |
| Juncus effusus | JUNUEFF | G | Gr | I |
| Juncus filiformis | JUNUFIL | D | Gr | I |
| Juncus squarrosus | JUNUSQU | C | Gr | aLI |
| Juncus tenuis | JUNUTEU | I | Gr | I |
| Juniperus communis | JUNIC.C | F | I | I |
| Knautia arvensis | KNAUARV | G | Fo | I |
| Koeleria pyramidata | KOELPYR | D | Gr | bLI |
| Lathyrus linifolius | LATYLIN | C | L | aLI |
| Lathyrus pratensis | LATYPRA | G | L | NI |
| Leontodon autumnalis | LEONAUT | G | Fo | I |
| Leontodon hispidus | LEONHIS | G | Fo | NI |
| Leucanthemum ircutianum | LEUNIRC | G | Fo | I |
| Lilium martagon | LILUMAR | I | Fo | I |
| Linum catharticum | LINUCAT | D | Fo | bLI |
| Lotus corniculatus | LOTUCOR | D | L | bLI |
| Lotus pendunculatus | LOTUPED | G | L | I |
| Lupinus polyphyllus | LUPIPOL | F | L | I |
| Luzula campestris | LUZUCAM | C | Gr | aLI |
| Luzula congesta | LUZUCON | C | Gr | I |
| Luzula luzuloides | LUZULUU | F | Gr | I |
| Luzula multiflora | LUZUMUL | C | Gr | I |
| Luzula sylvatica | LUZUS.Y | F | Gr | I |
| Lyopodium clavatum | LYCUCLV | C | Fo | aLI |
| Lysimachia vulgaris | LYSMVUL | I | Fo | I |
| Maianthemum bifolium | MAIABIF | F | Fo | aLI |
| Melampyrum pratensis | MELAPRA | F | Fo | aLI |
| Meum athamanticum | MEUMATH | D | Fo | aLI |
| Moehringia trinervia | MOEHTRI | F | Fo | NI |
| Molinia caerulea | MOLICAE | D | Gr | I |
| Myosotis nemorosa | MYOSNEM | G | Fo | I |
| Nardus stricta | NARUSTR | C | Gr | aLI |
| Pedicularis sylvatica | PEDISYL | C | Fo | aLI |
| Phyteuma nigrum | PHYENIG | G | Fo | I |
| Phyteuma orbiculare | PHYEO.O | D | Fo | bLI |
| Phyteuma spicatum | PHYESPI | I | Fo | I |
| Picea abies | PICEABI | F | I | I |
| Pimpinella major | PIMPM.M | G | Fo | NI |
| Pimpinella saxifraga | PIMPSAX | D | Fo | I |
| Pinus sylvestris | PINUSYL | F | I | I |
| Plantago lanceolata | PLAJLAN | G | Fo | I |
| Plantago major | PLAJM.M | I | Fo | NI |
| Platanthera bifolia | PLARBIF | I | Fo | I |
| Platanthera chlorantha | PLARCHL | D | Fo | I |
| Pleurozium schreberi | PLEZSCH | C | I | I |
| Poa chaixii | POA.CHA | I | Gr | I |
| Poa pratensis | POA.PRA | G | Gr | NI |
| Poa trivialis | POA.TRI | G | Gr | NI |
| Polygala serpyllifolia | POLGSER | C | Fo | aLI |
| Polygala vulgaris | POLGVUL | C | Fo | aLI |
| Populus tremula | POPUTRE | F | I | I |
| Potentilla erecta | POTEERE | D | Fo | I |
| Potentilla palustris | POTEPAL | D | Fo | aLI |
| Potentilla tabernaemontani | POTETAB | D | Fo | bLI |
| Primula elatior | PRIMELA | I | Fo | NI |
| Prunella grandiflora | PRUNGRA | D | Fo | bLI |
| Prunella vulgaris | PRUNVUL | G | Fo | I |
| Prunus avium | PRUUAVI | F | I | I |
| Prunus serotina | PRUUSER | F | I | I |
| Prunus spinosa | PRUUSPI | F | I | I |
| Pteridium aquilinum | PTERAQU | F | Fo | aLI |
| Quercus petraea | QUERPET | F | I | I |
| Quercus robur | QUERROB | F | I | I |
| Ranunculus acris | RANCACR | G | Fo | I |
| Ranunculus auricomus agg. | RANC.AU | I | Fo | I |
| Ranunculus bulbosus | RANCB.B | D | Fo | bLI |
| Ranunculus flammula | RANCFLA | D | Fo | aLI |
| Ranunculus nemorosus s.l. | RANC.PO | I | Fo | I |
| Ranunculus repens | RANCREP | I | Fo | NI |
| Rhinanthus minor | RHINMIN | D | Fo | I |
| Rhytidiadelphus squarrosus | RHYISQU | I | I | I |
| Roa canina | ROSACAN | F | I | I |
| Rubus fruticosus agg. | RUBU.FR | F | I | I |
| Rubus idaeus | RUBUIDA | F | I | NI |
| Rumex acetosa | RUMEACE | G | Fo | NI |
| Rumex acetosella | RUMEACT | D | Fo | aLI |
| Rumex obtusifolius | RUMEOBT | I | Fo | NI |
| Rumex sanguinea | RUMESAN | F | Fo | NI |
| Salix aurita | SALXAUR | F | I | I |
| Salix caprea | SALXCAP | F | I | NI |
| Salix repens | SALXREP | F | S | bLI |
| Salix x multinervis | SALX.MU | F | I | I |
| Sambucus nigra | SAMBNIG | F | I | NI |
| Sanguisorba minor | SANGM.M | D | Fo | bLI |
| Sanguisorba officinalis | SANGOFF | G | Fo | I |
| Saxifraga granulata | SAXIGRA | D | Fo | I |
| Senecio jacobaea | SENCJAC | G | Fo | I |
| Senecio sylvaticus | SENCSYL | F | Fo | NI |
| Serratula tinctoria | SERRTIN | D | Fo | bLI |
| Silene flos-cuculi | SILEFLO | G | Fo | I |
| Silene nutans | SILENUT | I | Fo | bLI |
| Silene otites | SILEOTI | D | Fo | bLI |
| Solidago virgaurea | SOLIV.V | I | Fo | I |
| Sorbus aucuparia | SORUAUC | F | I | I |
| Stellaria alsine | STELALS | I | Fo | I |
| Stellaria graminea | STELGRA | I | Fo | I |
| Succisa pratensis | SUCCPRA | D | Fo | I |
| Taraxacum sect. Ruderalia | TARA.RU | I | Fo | I |
| Tephroseris helenitis | TEPRHEL | D | Fo | I |
| Thesium pyrenaicum | THESPYR | D | Fo | I |
| Thymus pulegioides | THYUPUL | D | Fo | I |
| Tragopogon pratensis ssp. pratensis | TRAGP.P | G | Fo | NI |
| Trichophorum germanicum | TRIPC.G | D | Gr | aLI |
| Trientalis europaea | TRINEUR | F | Fo | aLI |
| Trifolium dubium | TRIFDUB | G | L | I |
| Trifolium medium | TRIFMED | I | L | bLI |
| Trifolium pratense | TRIFPRA | G | L | I |
| Trifolium repens | TRIFREP | G | L | NI |
| Trifolium spadiceum | TRIFSPA | D | Fo | aLI |
| Trisetum flavescens | TRIMFLA | G | Gr | I |
| Trollius europaeus | TROLEUR | G | Gr | I |
| Urtica dioica | URTIDIO | I | Fo | NI |
| Urtica urens | URTIURE | I | Fo | NI |
| Vaccinium myrtillus | VACIMYR | C | S | aLI |
| Vaccinium oxycoccus | VACIOXY | D | S | aLI |
| Vaccinium vitis-idaea | VACIVIT | C | S | aLI |
| Valeriana dioica | VALEDIO | G | Fo | I |
| Veronica chamaedrys | VEROCHA | I | Fo | I |
| Veronica herderifolia | VEROHED | I | Fo | NI |
| Veronica officinalis | VEROOFF | C | Fo | I |
| Viburnum opulus | VIBUOPU | F | I | NI |
| Vicia cracca | VICICRA | G | L | I |
| Vicia sepium | VICISEP | I | L | I |
| Viola canina | VIOLCAN | C | Fo | aLI |
| Viola palustris | VIOLPAL | D | Fo | aLI |
| Viola riviniana | VIOLRIV | F | Fo | I |
| Viola tricolor | VIOLT.T | G | Fo | I |

References

Ellenberg, H. 1992. *Zeigerwerte von Pflanzen in Mitteleuropa.* 2nd ed. Goltze, Göttingen.

Peppler-Lisbach, C. & Petersen, J. 2001. *Synopsis der Pflanzengesellschaften Deutschlands. Calluno-Ulicetea (G3), Teil 1: Nardetalia strictae - Borstgrasrasen,* Göttingen.
