## Supplementary Information S2 for "Long-term vegetation changes in species-rich *Nardus* grasslands of Central Germany indicate eutrophication, recovery from acidification and management change as main drivers"

**Appendix S2.** Detailed results of exact one-sample Wilcoxon tests on differences (∆) in deposition rates. Given are the Hodges-Lehmann estimator of location shift (Median ∆) and the 95% confidence interval of the estimator and the respective *P* values (Bauer 1972; Hothorn & Hornik 2015). Results for the overall dataset and the two study areas. Interaction FWB: RHN *P* value of two-sample exact Wilcoxon test ∆ v vs. study area.

| Deposition | Regional: overall | | | Local: FWB | | | Local: RHN | | | Interaction: FWB:RHN | *n* |
| --- | --- | --- | --- | --- | --- | --- | --- | --- | --- | --- | --- |
|  | Median ∆ | CI | *P* | Median ∆ | CI | *P* | Median ∆ | CI | *P* | *P* |  |
| ΔSOx | -13.15 | -14.31 – -11.81 | 0.000 | -9.32 | -10.31 – -8.50 | 0.000 | -17.37 | -17.97 – -16.72 | 0.000 | 0.000 | 40 |
| ΔNHy | 3.86 | 3.47 – 4.26 | 0.000 | 4.51 | 4.40 – 4.62 | 0.000 | 3.01 | 2.49 – 3.26 | 0.000 | 0.000 | 40 |
| ΔNOx | -0.55 | -0.89 – -0.28 | 0.000 | 0.16 | -0.15 – 0.41 | 0.284 | -1.32 | -1.41 – -1.22 | 0.000 | 0.000 | 40 |
| ΔNtot | 3.24 | 2.71 – 3.83 | 0.000 | 4.63 | 4.32 – 4.99 | 0.000 | 1.68 | 1.25 – 2.04 | 0.000 | 0.000 | 40 |
| ΔNHy:NOx | 0.42 | 0.40 – 0.44 | 0.000 | 0.42 | 0.41 – 0.44 | 0.000 | 0.41 | 0.35 – 0.45 | 0.000 | 0.213 | 40 |

Units: SOx: kg S ha^-1^ a^-1^ / Nhy, Nox, Ntot: kg N ha^-1^ a^-1^

References

Bauer, D.F. 1972. Constructing Confidence Sets Using Rank Statistics. *Journal of the American Statistical Association* 67(339): 687–690.

Hothorn, T. & Hornik, K. 2015. *exactRankTests. Exact Distributions for Rank and Permutation Tests. R package version 0.8-28*.
