## Supplementary Information S3 for "Long-term vegetation changes in species-rich *Nardus* grasslands of Central Germany indicate eutrophication, recovery from acidification and management change as main drivers"

**Appendix S3.** Detailed results of exact one-sample Wilcoxon tests on differences in soil variables, structural variables, Ellenberg indicator values, total species richness, species richness and cover of species groups. Given are the Hodges-Lehmann estimators of location shift (Median∆) and the 95% confidence interval of the estimators incl. the respective *P* values (Bauer 1972; Hothorn & Hornik 2015). Results for the overall dataset and the two study areas. FWB:RHN: *P* value of two-sample exact Wilcoxon test ∆v vs. study area; n no. of plots.

|  | Regional: overall | | | Local: FWB | | | Local: RHN | | | FWB:RHN | n |
| --- | --- | --- | --- | --- | --- | --- | --- | --- | --- | --- | --- |
|  | Med∆ | CI | *P* | Med∆ | CI | *P* | Med∆ | CI | *P* | *P* |  |
| **Soil parameters** |  |  |  |  |  |  |  |  |  |  |  |
| ∆pH | 0.25 | 0.15 – 0.35 | 0.000 | 0.33 | 0.2 – 0.4 | 0.000 | 0.1 | -0.02 – 0.32 | 0.183 | 0.022 | 94 |
| ∆CN | -0.94 | -1.37 – -0.45 | 0.000 | -0.95 | -1.55 – -0.3 | 0.007 | -0.94 | -1.54 – -0.23 | 0.011 | 0.978 | 80 |
| **Structural parameters** |  |  |  |  |  |  |  |  |  |  |  |
| ∆ cover shrub layer (%) | 2 | 1 – 21 | 0.000 | 2 | 1 – 21 | 0.000 |  |  |  | 0.012 | 87 |
| ∆ cover herb layer (%) | 0.5 | -2.5 – 4 | 0.768 | 4.5 | 0 – 7.5 | 0.021 | -5.5 | -10 – -1 | 0.009 | 0 | 87 |
| ∆ cover moss layer (%) | 19.75 | 11 – 30.5 | 0.000 | 19.25 | 10 – 28 | 0.000 | 25 | 2 – 49 | 0.030 | 0.656 | 87 |
| **Mean Ellenberg indicator values** |  |  |  |  |  |  |  |  |  |  |  |
| ∆ mean R (cover weighted) | 0.55 | 0.4 – 0.69 | 0.000 | 0.48 | 0.32 – 0.67 | 0.000 | 0.65 | 0.39 – 0.92 | 0.000 | 0.389 | 97 |
| ∆ mean R (p/a weighted) | 0.14 | 0.05 – 0.24 | 0.002 | 0.12 | 0.03 – 0.2 | 0.010 | 0.25 | -0.01 – 0.5 | 0.058 | 0.563 | 97 |
| ∆ mean N (cover weighted) | 0.33 | 0.22 – 0.44 | 0.000 | 0.33 | 0.22 – 0.44 | 0.000 | 0.33 | 0.1 – 0.56 | 0.009 | 0.845 | 97 |
| ∆ mean N (p/a weighted) | 0.32 | 0.25 – 0.39 | 0.000 | 0.29 | 0.21 – 0.37 | 0.000 | 0.38 | 0.23 – 0.53 | 0.058 | 0.275 | 97 |
| **Species richness** |  |  |  |  |  |  |  |  |  |  |  |
| Δ total species richness | -0.25 | -2.5 – 2.5 | 0.958 | 0.75 | -1.5 – 3.5 | 0.358 | -2.25 | -7.5 – 4 | 0.458 | 0.168 | 97 |
| **Sociological groups** |  |  |  |  |  |  |  |  |  |  |  |
| Δ richness character species | -1.75 | -2.5 – -1 | 0.000 | -1.25 | -2.5 – -0.5 | 0 | -2.25 | -4 – 0 | 0.013 | 0.446 | 97 |
| Δ cover character species | -23 | -34 – -12 | 0.000 | -14 | -26.5 – -1.5 | 0.028 | -40.5 | -61.5 – -19 | 0.000 | 0.032 | 97 |
| Δ richness other low prod. grassl. spec. | -2.25 | -3 – -1 | 0.000 | -1.25 | -2.5 – -0.5 | 0.001 | -2.75 | -5 – 0 | 0.025 | 0.104 | 97 |
| Δ cover other low prod. grassl. spec. | -7 | -12 – -1.5 | 0.016 | 2.47 | -4.24 – 9 | 0.467 | -21.5 | -29 – -14 | 0.000 | 0.000 | 97 |
| Δ richness grassland species | 1.75 | 0.5 – 3 | 0.005 | 1.75 | 0.5 – 3 | 0.000 | 1.75 | -2 – 5 | 0.229 | 0.277 | 97 |
| Δ cover grassland species | 10.5 | 4 – 17 | 0.002 | 19 | 11 – 26 | 0.000 | -2.5 | -14 – 6.5 | 0.509 | 0.001 | 97 |
| Δ richness fallow species | 0.75 | 0.5 – 2 | 0.000 | 1.25 | 0.5 – 2 | 0.000 | 0.25 | -1.5 – 1 | 0.650 | 0.013 | 97 |
| Δ cover fallow species | 1.5 | 0 – 3 | 0.013 | 3 | 1.5 – 5.5 | 0.000 | -1.25 | -5 – 1 | 0.438 | 0.003 | 97 |
| **Ecological groups** |  |  |  |  |  |  |  |  |  |  |  |
| Δ richness nutrient ind. | 1.25 | 0.5 – 2 | 0.000 | 0.75 | 0.5 – 2 | 0.000 | 1.75 | 0 – 3 | 0.011 | 0.95 | 97 |
| Δ cover nutrient ind. | 2.5 | 1 – 3.5 | 0.000 | 3 | 2 – 5 | 0.000 | 1 | -1.5 – 3.5 | 0.397 | 0.116 | 97 |
| Δ richness basiphytic low nutrient ind. | -1.25 | -1.5 – -1 | 0.000 | -1.25 | -1.5 – -1 | 0.000 | -1.25 | -2.5 – 0 | 0.039 | 0.738 | 97 |
| Δ cover basiphytic low nutrient ind. | -2.5 | -4 – -1.5 | 0.000 | -2 | -3 – -0.5 | 0.014 | -3.5 | -6.5 – -1.5 | 0.004 | 0.125 | 97 |
| Δ richness acidophytic low nutrient ind. | -1.25 | -2.5 – -0.5 | 0.000 | -1.25 | -2 – 0 | 0.002 | -2.25 | -3.5 – 0 | 0.018 | 0.205 | 97 |
| Δ cover acidophytic low nutrient ind. | -25 | -33.5 – -16 | 0.000 | -16 | -27.5 – -6 | 0.003 | -38 | -53.5 – -24.5 | 0.000 | 0.009 | 97 |
| **Functional groups** |  |  |  |  |  |  |  |  |  |  |  |
| ∆ proportional richness graminoids | -0.02 | -0.04 – 0 | 0.054 | -0.01 | -0.04 – 0.01 | 0.286 | -0.03 | -0.08 – 0 | 0.062 | 0.339 | 97 |
| ∆ proportional cover graminoids | -0.05 | -0.08 – -0.01 | 0.008 | -0.04 | -0.08 – 0 | 0.033 | -0.05 | -0.12 – 0.01 | 0.097 | 0.748 | 97 |
| ∆ proportional richness forbs | 0.04 | 0.02 – 0.07 | 0.001 | 0.04 | 0.01 – 0.07 | 0.022 | 0.06 | 0.01 – 0.12 | 0.014 | 0.397 | 97 |
| ∆ proportional cover forbs | 0.05 | 0.02 – 0.09 | 0.001 | 0.04 | 0 – 0.08 | 0.044 | 0.08 | 0.02 – 0.15 | 0.009 | 0.196 | 97 |
| ∆ graminoid:forb ratio (p/a) | -0.11 | -0.21 – -0.01 | 0.028 | -0.08 | -0.21 – 0.05 | 0.258 | -0.15 | -0.32 – -0.01 | 0.027 | 0.313 | 97 |
| ∆ graminoid:forb ratio (cover) | -0.27 | -0.55 – -0.02 | 0.035 | -0.23 | -0.57 – 0.07 | 0.122 | -0.33 | -0.94 – 0.16 | 0.158 | 0.72 | 97 |

References

Bauer, D.F. 1972. Constructing Confidence Sets Using Rank Statistics. *Journal of the American Statistical Association* 67(339): 687–690.

Hothorn, T. & Hornik, K. 2015. *exactRankTests. Exact Distributions for Rank and Permutation Tests. R package version 0.8-28*.
