## Supplementary Information S4 for "Long-term vegetation changes in species-rich *Nardus* grasslands of Central Germany indicate eutrophication, recovery from acidification and management change as main drivers"

**Appendix S4.** Detailed results of exact one-sample Wilcoxon tests on differences (∆) in species frequency and cover. Given are overall frequencies in the data set (frequ.), overall mean cover (%). number of plots with changes of the respective species (no. ch.), mean differences (∆) in species frequency and cover (%) and the P-values of the one-sample exact Wilcoxon tests on ∆frequency (p.adj.fr) and ∆cover (p.adj.cover). P values were adjusted for multiple testing using the false discovery rate method of Benjamini & Hochberg (1995). FWB:RHN: significant differences (two-sample Wilcoxon test) in cover (cov) or frequency between study areas (FWB): only occurring in FWB plots, (RHN): only occurring in RHN plots. sG. Sociological species group; EIV R Ellenberg R indicator value, EIV N Ellenberg N indicator value (Ellenberg 1992); mean: mean R and N values of decreasing and increasing species.

|  | **Species** | **sG** | **EIV R** | **EIV R** | **frequ.** | **mean cover (%)** | **no. ch** | **∆freq. (%)** | **∆cov. (%)** | **p.adj. fr.** | **p.adj. cov.** | **FWB: RHN** |
| --- | --- | --- | --- | --- | --- | --- | --- | --- | --- | --- | --- | --- |
| decrease in | *Danthonia decumbens* | C | 3 | 2 | 113 | 2.12 | 41 | -29.90 | -0.54 | 0.000 | 0.010 |  |
| cover and | *Arnica montana* | C | 3 | 2 | 62 | 2.27 | 26 | -24.74 | -0.95 | 0.000 | 0.004 |  |
| frequency | *Festuca ovina* agg. | D |  |  | 144 | 5.01 | 32 | -24.74 | -2.71 | 0.000 | 0.011 |  |
|  | *Campanula rotundifolia* | D |  | 2 | 96 | 1.31 | 24 | -18.56 | -0.57 | 0.004 | 0.000 | cov |
|  | *Briza media* | D |  | 2 | 32 | 0.37 | 24 | -18.56 | -0.41 | 0.004 | 0.015 |  |
|  | *Thymus pulegioides* | D |  |  | 37 | 0.47 | 23 | -17.53 | -0.43 | 0.006 | 0.013 |  |
|  | *Polytrichum commune* s.l. |  |  |  | 33 | 1.77 | 21 | -15.46 | -1.88 | 0.013 | 0.015 |  |
|  | *Nardus stricta* | C | 2 | 2 | 173 | 23.58 | 17 | -13.40 | -14.01 | 0.016 | 0.000 |  |
|  | *Plagiomnium affine* |  | 5 |  | 21 | 0.63 | 15 | -11.34 | -1.06 | 0.037 | 0.015 | (FWB) |
|  | *Solidago virgaurea* |  |  | 4 | 15 | 0.18 | 9 | -9.28 | -0.21 | 0.023 | 0.023 |  |
| decrease in | *Calluna vulgaris* | C | 1 | 1 | 73 | 2.27 | 29 | -23.71 | -1.49 | 0.000 | 0.051 |  |
| frequency only | *Carex panicea* | D |  | 4 | 42 | 1.06 | 16 | -12.37 | -0.39 | 0.023 | 0.051 |  |
|  | *Vaccinium myrtillus* | C | 2 | 3 | 110 | 6.27 | 34 | -26.80 | -0.62 | 0.000 | 0.061 |  |
|  | *Hieracium pilosella* | D |  | 2 | 69 | 1.34 | 27 | -17.53 | -0.18 | 0.013 | 0.256 |  |
| decrease in | *Deschampsia flexuosa* | C | 2 | 3 | 129 | 8.36 | 25 | -7.22 | -6.79 | 0.459 | 0.006 |  |
| cover only | *Cirsium acaule* | D | 8 | 2 | 20 | 0.27 | 14 | -8.25 | -0.41 | 0.192 | 0.015 |  |
|  | *Vaccinium vitis-idaea* | C | 2 | 1 | 19 | 0.30 | 9 | -7.22 | -0.31 | 0.144 | 0.031 |  |
|  | *Trollius europaeus* | G | 6 | 5 | 22 | 0.22 | 12 | -6.19 | -0.21 | 0.356 | 0.038 | (RHN) |
|  | *Antennaria dioica* | D | 3 | 2 | 11 | 0.24 | 11 | -7.22 | -0.42 | 0.196 | 0.047 |  |
|  |  | mean: | 3.4 | 2.5 |  |  |  |  |  |  |  |  |
| increase in | *Picea abies* juv. | B |  |  | 11 | 0.10 | 9 | 9.28 | 0.14 | 0.023 | 0.023 |  |
| cover and | *Platanthera bifolia* |  | 7 |  | 9 | 0.06 | 9 | 9.28 | 0.11 | 0.023 | 0.023 |  |
| frequency | *Ceratodon purpureus* |  |  |  | 10 | 0.19 | 10 | 10.31 | 0.37 | 0.014 | 0.015 | (FWB) |
|  | *Geranium sylvaticum* | G | 6 | 7 | 12 | 0.21 | 12 | 10.31 | 0.37 | 0.034 | 0.035 | (RHN) |
|  | *Vicia cracca* | G |  |  | 26 | 0.30 | 16 | 12.37 | 0.25 | 0.023 | 0.022 |  |
|  | *Hypocharis radicata* | D | 4 | 3 | 22 | 0.29 | 14 | 12.37 | 0.29 | 0.014 | 0.029 |  |
|  | *Rhinanthus minor* | D |  | 3 | 49 | 0.86 | 21 | 13.40 | 0.54 | 0.037 | 0.022 |  |
|  | *Trifolium pratense* | G |  |  | 32 | 0.71 | 22 | 16.49 | 1.05 | 0.009 | 0.002 |  |
|  | *Trifolium repens* | G | 6 | 6 | 29 | 0.77 | 21 | 17.53 | 1.25 | 0.004 | 0.001 |  |
|  | *Stellaria graminea* |  | 4 | 3 | 53 | 0.59 | 25 | 17.53 | 0.38 | 0.009 | 0.015 |  |
|  | *Taraxacum* sect. Ruderalia |  |  | 7 | 22 | 0.20 | 18 | 18.56 | 0.31 | 0.000 | 0.000 |  |
|  | *Holcus lanatus* | G |  | 5 | 76 | 1.09 | 26 | 18.56 | 0.66 | 0.006 | 0.005 |  |
|  | *Veronica chamaedrys* |  |  |  | 75 | 1.38 | 31 | 27.84 | 1.52 | 0.000 | 0.000 |  |
|  | *Rhytidiadelphus squarrosus* |  | 5 |  | 107 | 14.67 | 39 | 36.08 | 22.70 | 0.000 | 0.000 |  |
|  |  | mean: | 5.3 | 4.9 |  |  |  |  |  |  |  |  |

References

Benjamini, Y. & Hochberg, Y. 1995. Controlling the False Discovery Rate: A Practical and Powerful Approach to Multiple Testing. *Journal of the Royal Statistical Society: Series B (Methodological)* 57(1): 289–300.

Ellenberg, H. 1992. *Zeigerwerte von Pflanzen in Mitteleuropa.* 2nd ed. Goltze, Göttingen.
