## Supplementary Information S5 for "Long-term vegetation changes in species-rich *Nardus* grasslands of Central Germany indicate eutrophication, recovery from acidification and management change as main drivers"

**Appendix S5.** Detailed results of linear regression models of changes in mean indicator values, species richness and species groups on environmental drivers, management change and study area. Given are explained variance (R²), *P*-value of the overall model, regression coefficients and respective *P* values.

ΔM management change: FM fallow to managed; MM continuously managed; reference category: xF fallow at resurvey

SA: study area: RHN Rhön; reference category FWB Fulda-Werra-Bergland

| **Variable** | **R^2^** | ***P*** | **Interc.** | ***P*** | **Δ pH** | ***P*** | **Δ CN** | ***P*** | **ΔM: FM** | ***P*** | **ΔM: MM** | ***P*** | **SA: RHN** | ***P*** |
| --- | --- | --- | --- | --- | --- | --- | --- | --- | --- | --- | --- | --- | --- | --- |
| **Mean indicator values** |  |  |  |  |  |  |  |  |  |  |  |  |  |  |
| Δ mean R value (cover-weighted) | 0.25 | 0.000 | -0.31 | 0.009 | 1.13 | 0.000 |  |  |  |  |  |  |  |  |
| Δ mean R value (p/a) | 0.24 | 0.000 | 0.01 | 0.834 | 0.54 | 0.000 |  |  |  |  |  |  |  |  |
| Δ mean N value (cover-weighted) |  | n.s. |  |  |  |  |  |  |  |  |  |  |  |  |
| Δ mean N value (p/a) |  | n.s. |  |  |  |  |  |  |  |  |  |  |  |  |
| **Species richness** |  |  |  |  |  |  |  |  |  |  |  |  |  |  |
| ∆ total richness | 0.20 | 0.000 | -0.27 | 0.024 | 1.01 | 0.000 |  |  |  |  |  |  |  |  |
| **Sociological species groups** |  |  |  |  |  |  |  |  |  |  |  |  |  |  |
| Δ richness character species |  |  |  |  |  |  |  |  |  |  |  |  |  |  |
| Δ cover character species | 0.06 | 0.026 | 0.19 | 0.177 |  |  |  |  |  |  |  |  | -0.51 | 0.026 |
| Δ richness other low-prod. grassl. sp. | 0.29 | 0.000 | 0.01 | 0.919 | 0.95 | 0.000 |  |  |  |  |  |  | -0.75 | 0.000 |
| Δ cover other low prod. grassland sp. | 0.22 | 0.000 | -0.29 | 0.015 | 1.07 | 0.000 |  |  |  |  |  |  |  |  |
| Δ richness agricultural grassland sp. | 0.15 | 0.000 | -0.24 | 0.055 | 0.88 | 0.000 |  |  |  |  |  |  |  |  |
| Δ cover agricultural grassland sp. | 0.13 | 0.001 | -0.22 | 0.074 | 0.83 | 0.001 |  |  |  |  |  |  |  |  |
| Δ richness fallow species | 0.12 | 0.007 | 0.64 | 0.029 |  |  |  |  | -0.52 | 0.117 | -1.02 | 0.003 |  |  |
| Δ cover fallow species | 0.21 | 0.000 | 1.01 | 0.000 |  |  |  |  | -0.97 | 0.002 | -1.41 | 0.000 |  |  |
| **Ecological species groups** |  |  |  |  |  |  |  |  |  |  |  |  |  |  |
| Δ richness nutrient indicators | 0.08 | 0.013 | -0.17 | 0.183 | 0.63 | 0.013 |  |  |  |  |  |  |  |  |
| Δ cover nutrient indicators | 0.08 | 0.013 | -0.17 | 0.183 | 0.63 | 0.013 |  |  |  |  |  |  |  |  |
| Δ richness basiphytic low nutrient ind. | 0.21 | 0.000 | -0.28 | 0.018 | 1.05 | 0.000 |  |  |  |  |  |  |  |  |
| Δ cover basiphytic low nutrient ind. | 0.20 | 0.000 | -0.27 | 0.024 | 1.01 | 0.000 |  |  |  |  |  |  |  |  |
| Δ richness acidophytic low nutrient ind. |  |  |  |  |  |  |  |  |  |  |  |  |  |  |
| Δ cover acidophytic low nutrient ind. | 0.08 | 0.011 | 0.21 | 0.120 |  |  |  |  |  |  |  |  | -0.59 | 0.011 |
| **Functional species groups** |  |  |  |  |  |  |  |  |  |  |  |  |  |  |
| Δrichness graminoids | 0.14 | 0.003 | 0.08 | 0.515 | -0.70 | 0.005 | -0.13 | 0.020 |  |  |  |  |  |  |
| Δcover graminoids | 0.16 | 0.001 | 0.06 | 0.617 | -0.69 | 0.005 | -0.15 | 0.006 |  |  |  |  |  |  |
| Δrichness forbs | 0.12 | 0.002 | -0.21 | 0.087 | 0.80 | 0.002 |  |  |  |  |  |  |  |  |
| Δcover forbs | 0.15 | 0.000 | -0.24 | 0.055 | 0.88 | 0.000 |  |  |  |  |  |  |  |  |
| ΔGr:Fo ratio (richness) |  | n.s. |  |  |  |  |  |  |  |  |  |  |  |  |
| ΔGr:Fo ratio (cover) | 0.08 | 0.013 | -0.17 | 0.182 | 0.63 | 0.013 |  |  |  |  |  |  |  |  |
