## Supplementary Information S6 for "Long-term vegetation changes in species-rich *Nardus* grasslands of Central Germany indicate eutrophication, recovery from acidification and management change as main drivers"

**Appendix S6. (a)** RDA of species differences (p/a species data) with ΔpH and ΔM (FF vs. FM and MM) as constraining variables. Species data were controlled for study area (conditioning variable). For detailed results see Appendix S6b.
Blue arrows: constraining variables; FF1: fallow at resurvey, FF0: managed at resurvey; Δ pH change in soil pH; Green arrows: supplementary variables (correlations).

Abbrev.: d. ∆ of p/a weighted data; mR mean p/a R value; mN mean p/a N value; tRi total species richness; C character species; D other low-productive grassland species; G agricultural grassland species; F fallow species; aLI acidophytic low nutrient indicators; bLI basiphytic low nutrient indicators; NI nutrient indicators; pFo proportion of forbs; pGr proportion of graminoids; .1 tree layer; .2 shrub layer; .3 herb layer; .4 moss layer.
Inclusion rules for species: no. of plots with changes > 4 & *P*-value of multiple linear regression of p/a values on axis 1+2 scores < 0.05.

-1.0

-0.5

0.0

0.5

1.0

1.5

2.0

-1.5

-1.0

-0.5

0.0

0.5

1.0

RDA1

RDA2

ACHIMIL.3

ALCH.VU.3

ALCHGLU.3

ANTXODO.3

ARRHELA.3

BRIZMED.3

CARDPRA.3

CARENIG.3

CAREPAN.3

CAREPIU.3

CIRSPAL.3

CYNSCRI.3

DANTDEC.3

DICUBON.4

GALOTET.3

GALUBOR.3

GALUPUM.3

GALUSAX.3

GALUVEU.3

GERISYL.3

HELTPRA.3

HIERLAT.3

HIERPIO.3

HOLCLAN.3

HYPCRAD.3

HYPEMAC.3

KNAUARV.3

LATYPRA.3

LEONHIS.3

LEUNIRC.3

LOTUCOR.3

LUZUCAM.3

LUZUMUL.3

NARUSTR.3

PIMPSAX.3

PLAJLAN.3

PLEZSCH.4

POLGVUL.3

POLZFOR.4

QUERROB.2

RANCACR.3

RANC.PO.3

RHYISQU.4

RUBUIDA.2

RUMEACE.3

SUCCPRA.3

TARA.RU.3

THYUPUL.3

TRIFMED.3

TRIFPRA.3

TRIFREP.3

VACIMYR.3

VACIVIT.3

VEROCHA.3

VEROOFF.3

VIOLCAN.3

VIOLT.T.3

FF0

FF1

d.ph

d.tR

d.C

d.D

d.G

d.F

d.mRpa

d.mNpa

d.pGr

d.pFo

d.NI

d.bLI

d.aLI

**Appendix S6. (b)** Detailed results of redundancy analyses.

RDA-1a: RDA of species differences (cover-weighted species data) with Δ pH and Δ M (FF vs. FM and MM) as constraining variables. Conditioning variable: Study area.

RDA-1b: RDA of species differences (p/a species data) with Δ pH and Δ M (FF vs. FM and MM) as constraining variables. Conditioning variable: Study area.

RDA-2: RDA of species differences (p/a-data) with Δ pH and Δ NHy as constraining variables. Conditioning variables: Study area, altitude and management

Given are % explained variance of conditioning and constraining variables and results of permutations tests (1000 permutations) for significance of axes and constraining variables. Constraining variables were assessed sequentially following the order in column “Constraining variables”.

| **Ordination** | **Conditioning variables** | **Constraining variables** | **no. obs.** | **% var. cond.** | **% var. constr.** | **% var. axis 1** | **% var. axis 2** | ***P* axis 1** | ***P* axis 2** | ***P* ∆pH** | ***P* FF** | ***P* ∆NHy** |
| --- | --- | --- | --- | --- | --- | --- | --- | --- | --- | --- | --- | --- |
| RDA-1a | SA | Δ pH+ Δ M (FF) | 80 | 3.00 | 4.81 | 2.95 | 2.01 | 0.002 | 0.018 | 0.002 | 0.016 |  |
| RDA-1b | SA | Δ pH+ Δ M (FF) | 80 | 1.94 | 4.45 | 2.97 | 1.59 | 0.001 | 0.063 | 0.001 | 0.044 |  |
| RDA-2 | SA + alt + ∆ M | Δ pH+ Δ NHy | 40 | 8.17 | 8.23 | 5.19 | 3.77 | 0.003 | 0.029 | 0.002 |  | 0.008 |
